# Somatostatin released by single O-LM interneurons slowly inhibits intrinsic neuronal excitability in a paracrine and autocrine manner

**DOI:** 10.1101/2025.08.31.673310

**Authors:** Maria Laura Musella, Malika Sammari, Chaima Ajal, Anaïs Le Cor, Michaël Russier, Dominique Debanne, Salvatore Incontro

## Abstract

Somatostatin (SST) is a neuropeptide known to inhibit both neuronal excitability and excitatory synaptic transmission in principal neurons. While SST is expressed in some GABAergic interneurons such as *Oriens-Lacunosum Moleculare* (O-LM) cells, the precise conditions governing SST release remain poorly defined. We show here that, in the presence of GABA_A_ receptor antagonist, single O-LM interneurons of CA1 are able to inhibit pyramidal neuron excitability when O-LM cells fire at frequencies above 20 Hz. This inhibition is suppressed by the SST receptor antagonist cyclo-SST and is absent when the O-LM interneuron is located more than 150 µm from the pyramidal cell. Likewise, a single O-LM cell was also able to inhibit the intrinsic excitability of another O-LM cell, provided the two cells are in proximity. Exogeneous application of SST transiently inhibits excitatory synaptic transmission and intrinsic excitability in O-LM interneurons through SST1 and SST2, respectively. Notably, this transient inhibition became sustained when clathrin-dependent endocytosis of SST receptors was blocked. Blocking of SST receptors reduced NMDA receptor-mediated responses, prevented induction of both synaptic and intrinsic potentiation in pyramidal neurons and reduced the coherence of theta oscillations. These findings reveal that SST released by O-LM interneurons constitutes an additional inhibitory mechanism, modulating both synaptic excitation and intrinsic excitability beyond the transient inhibition mediated by GABA release.

## Introduction

Oriens-Lacunosum Moleculare (O-LM) interneurons play a crucial role in brain activity (Fernández-Arroyo et al., 2025). These interneurons contribute to oscillatory dynamics (Klausberger et al., 2003; Gloveli et al., 2005; Goldin et al., 2007; Tort et al., 2007; Mikulovic et al., 2018), express synaptic plasticity and intrinsic plasticity (Incontro et al., 2021; Sammari et al., 2022), are associated with memory formation (Schmid et al., 2016; Siwani et al., 2018; Udakis et al., 2020; Taxidis et al., 2025), and participate to complex behaviors (Katona et al., 2014; Hainmueller et al., 2024). Because they directly target distal apical dendrites of CA1 pyramidal neurons and inhibit interneurons targeting *stratum radiatum* dendrites, O-LM interneurons are able to discriminate CA3 inputs mainly targeting the *stratum radiatum* from entorhinal inputs targeting the *lacunosum moleculare* (Leão et al., 2012). These interneurons fire during theta episodes at frequencies ranging between 5 and 50 Hz (Klausberger et al., 2003).

Somatostatin (SST) is a neuropeptide expressed in several hippocampal GABAergic interneuron subtypes including O-LM and bistratified interneurons (Müller and Remy, 2014). SST inhibits excitatory synaptic transmission (Tallent and Siggins, 1997) and neuronal excitability in principal neurons (Moore et al., 1988). SST exerts its effect via five G protein-coupled receptors (SST1-SST5) which are differentially distributed in the brain, pre- and post-synaptically (Csaba and Dournaud, 2023). In hippocampal CA1 pyramidal neurons, SST1 mediates presynaptic inhibition of excitatory transmission (Cammalleri et al., 2009) while SST2 reduces postsynaptic intrinsic excitability (Dournaud et al., 1996). Inhibition of neuronal excitability results from the activation of both Kv7 channel-mediated M-current (Moore et al., 1988; Qiu et al., 2008) through arachidonic acid and its metabolites (Schweitzer et al., 1990, 1993), as well as the enhancement of leak K^+^ current (Schweitzer et al., 1998). O-LM interneurons thus harbor two potent inhibitory transmitters: GABA and SST. While the dynamics of GABA release from O-LM onto CA1 pyramidal cell has been extensively studied (Maccaferri et al., 2000), the release of SST by GABAergic interneuron is surprisingly much less defined.

Using dual recording from an O-LM interneuron and either a CA1 pyramidal cell or another neighboring O-LM interneuron, we show in this study that SST released from a single O-LM interneuron can inhibit both pyramidal and O-LM excitability when the neurons are close to each other (<∼100-150 µm) and when the firing in the O-LM exceeds 20 Hz. Constant SST application inhibits transiently both excitatory synaptic transmission and intrinsic neuronal excitability in O-LM interneurons. The transient nature of SST-mediated inhibition is due to the clathrin-dependent endocytosis of SST1 at the presynaptic side and SST2 at the postsynaptic side. Thus, SST released by O-LM interneurons provides a complementary inhibitory mechanism that reduces synaptic excitation and intrinsic excitability, prolonging the brief, phasic inhibition mediated by GABA into a slower and sustained form of circuit control.

## Results

### Paracrine inhibition of SST released by a single O-LM interneuron

Dual whole-cell recordings from adjacent O-LM interneuron and pyramidal cell were obtained in the CA1 area from acute slices of rat dorsal hippocampus in order to test the effect of SST release by the O-LM on pyramidal cell firing. GABA_A_ channels were blocked with picrotoxin (100 µM). Pyramidal cell firing was evoked by current pulses at three time points (**Figure 1A**) while O-LM interneurons were subject to four conditions of stimulation: i) no current, ii) subthreshold depolarization, iii) firing below 20 Hz and iv) firing above 20 Hz (**Figure 1A**). Only firing above 20 Hz in O-LM cells inhibited pyramidal cell firing (**Figure 1B**). This firing frequency is commonly observed in O-LM interneurons during network activity (**Figure S1**). This inhibition was ∼10% at 2.5 (Wilcoxon test, p<0.01) and 10 s (Wilcoxon test, p<0.05) but absent at 20 s (Wilcoxon test, ns) post O-LM firing (**Figure 1B**). Firing below 20 Hz had no effect on pyramidal cell excitability (**Figure S2**) and as expected, pyramidal cell firing remained unchanged when no current injection and subthreshold depolarization was induced in the O-LM (**Figure 1B**). Importantly, in the presence of the somatostatin receptor antagonist, cyclo-somatostatin (cyclo-SST, 2 µM), neuronal excitability remained unchanged in pyramidal neurons upon O-LM interneuron firing at a frequency larger than 20 Hz (**Figure 1B**). These results strongly suggest that SST released by O-LM interneuron inhibits CA1 pyramidal cells.

**Figure 1.**
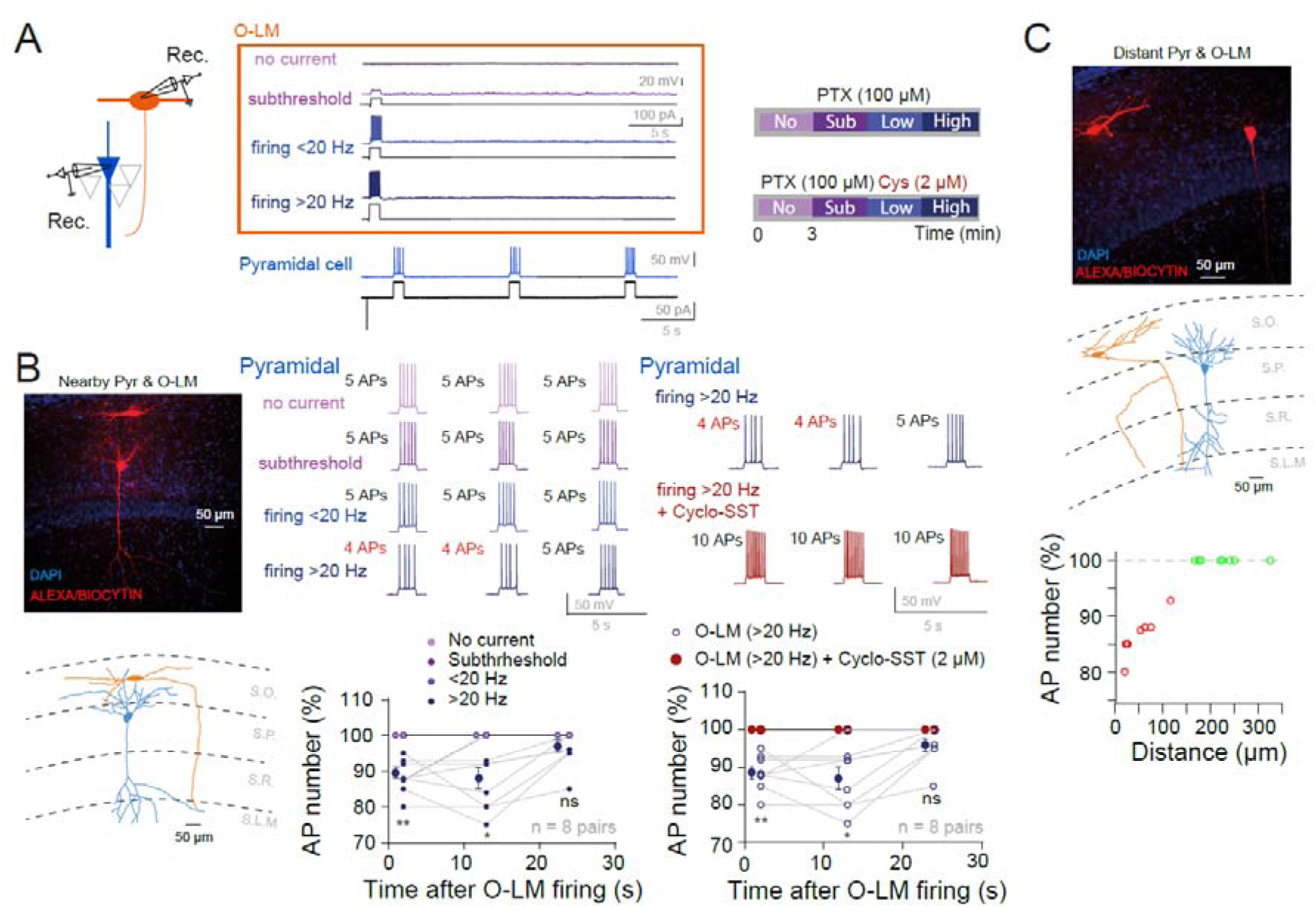
SST released by single O-LM interneuron inhibits CA1 pyramidal cell excitability. A. Recording configuration and stimulation protocol. B. Inhibition of pyramidal cells by nearby O-LM interneurons. Left, image and reconstruction of pyramidal and O-LM neurons labelled with biocytin. Middle, pyramidal cell excitability is reduced when the O-LM interneuron fire at a frequency larger than 20 Hz. Top, voltage traces in a pyramidal neuron at different times and mode of firing. Bottom, quantitative data. **, p<0.01; *, p<0.05, ns, not significant. Right, cyclo-SST suppresses the O-LM firing mediated inhibition. Top, inhibition traces. Bottom, quantitative data showing the total suppression of SST-mediated inhibition. C. Lack of inhibition by distant O-LM interneurons. Top, image and reconstruction of distant pyramidal and O-LM neurons labelled with biocytine. Bottom, normalized AP number evoked at 0.2 s as a function of the distance between the two neurons. Note that at distance >150 µM, no inhibition is observed.

We then increased the distance between the recorded pyramidal cell and the O-LM interneuron (**Figure 1C**). At distance larger than 150 µm, spiking in the O-LM did not inhibit pyramidal cell firing (**Figure 1C**). Notably, cyclo-SST increased pyramidal neuron excitability (**Figure S3**), indicating that postsynaptic SST receptors are constitutively activated by background SST. These results indicate that SST released by a single O-LM interneuron can inhibit CA1 pyramidal cells via a paracrine signaling within a ∼150 µm radius.

We next examined whether a single O-LM interneuron could inhibit the firing of adjacent O-LM cells (**Figure 2**). Using the same picrotoxin-base disinhibition protocol, dual recordings were obtained from two adjacent O-LM interneurons. Firing was evoked in one O-LM interneuron with current pulses of constant amplitude at three time points while four modes of stimulation were applied at the other O-LM cell that was tested under the same four paradigms described above (**Figure 2A**). Again, only O-LM firing above 20 Hz inhibited the firing of the neighboring O-LM interneuron (**Figure 2B**). The inhibition was ∼15% at 2.5 (Wilcoxon test, p<0.05) and 10 s (Wilcoxon test, p<0.01) but almost disappeared at 20 s (Wilcoxon test, ns) post O-LM firing (**Figure 2B**). Lower firing frequencies (< 20 Hz) had no effect (**Figure S4**) and, as expected, cell firing remained unchanged when no current injection and subthreshold depolarization was induced in the O-LM (**Figure 2B**). Importantly, this inhibition was abolished by cyclo-SST (2 µM) (**Figure 2B**), confirming an SST-mediated mechanism. These results strongly suggest that SST released by an O-LM interneuron inhibits other O-LM cells.

**Figure 2.**
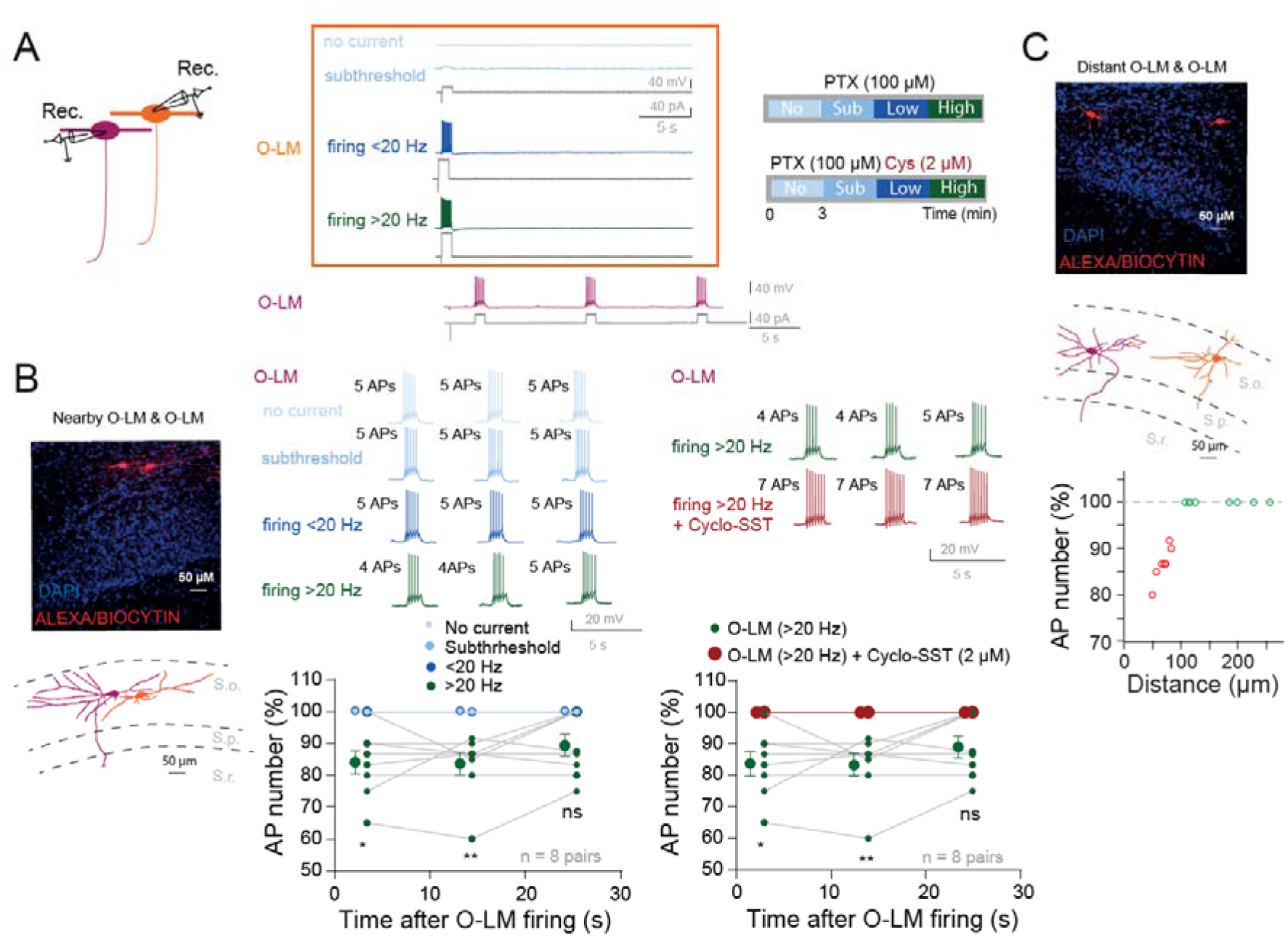
SST released by one O-LM inhibits neighbouring O-LM interneurons. A. Left, recording configuration of pairs of O-LM interneurons. Middle, experimental protocol. Right, time line of the protocol. B. Left, morphology of two proximal O-LM interneurons. Top, confocal images. Bottom, image and reconstruction of two distant O-LM interneurons labelled with biocytin. Middle, O-LM cell excitability is reduced when the O-LM interneuron fire at a frequency larger than 20 Hz. Right, blockade of SST receptors with cyclo-SST suppress the inhibition of O-LM cell firing. C. Top, confocal images of two distant O-LM interneurons. Middle, reconstruction of the two O-LM cells. Bottom, inhibition as a function of the distance between the two O-LM cells.

Increasing the intersomatic distance between O-LM pairs revealed that inhibition was absent when cells were separated by more than 100 µm (**Figure 2C**). As for pyramidal neurons, cyclo-SST increased O-LM cell excitability (**Figure S5**), indicating that SST receptors are constitutively activated by ambient SST.

These results demonstrate that SST released from a single O-LM interneuron exerts paracrine inhibition within a spatial range of ∼150 µm for pyramidal cells and ∼100 µm for other O-LM cells.

### Autocrine inhibition of SST released by a single O-LM interneuron

We next investigate whether SST released by a single O-LM interneuron could self-inhibit its own firing. To check this, O-LM interneurons were recorded in acute slices of rat dorsal hippocampus. O-LM excitability was tested before and after a depolarizing pulse of current eliciting firing below or above 20 Hz (**Figure 3A**). Low firing frequency (< 20 Hz) had no effect on excitability (Wilcoxon test, p>0.1; **Figure 3B**) whereas high firing frequency (> 20 Hz) produced a clear inhibition in O-LM firing (Wilcoxon test, p<0.01; **Figure 3B**). This inhibition was abolished by either the broad-spectrum SST receptor antagonist cyclo-SST (2 µM) or the SST2-selective antagonist CYN154806 (1 µM) (Wilcoxon test, ns; **Figure 3B** and Wilcoxon test, ns; **Figure 3C**). Bath application of either antagonists increased O-LM excitability (**Figure S6**), indicating tonic activation of SST2 receptors.

**Figure 3.**
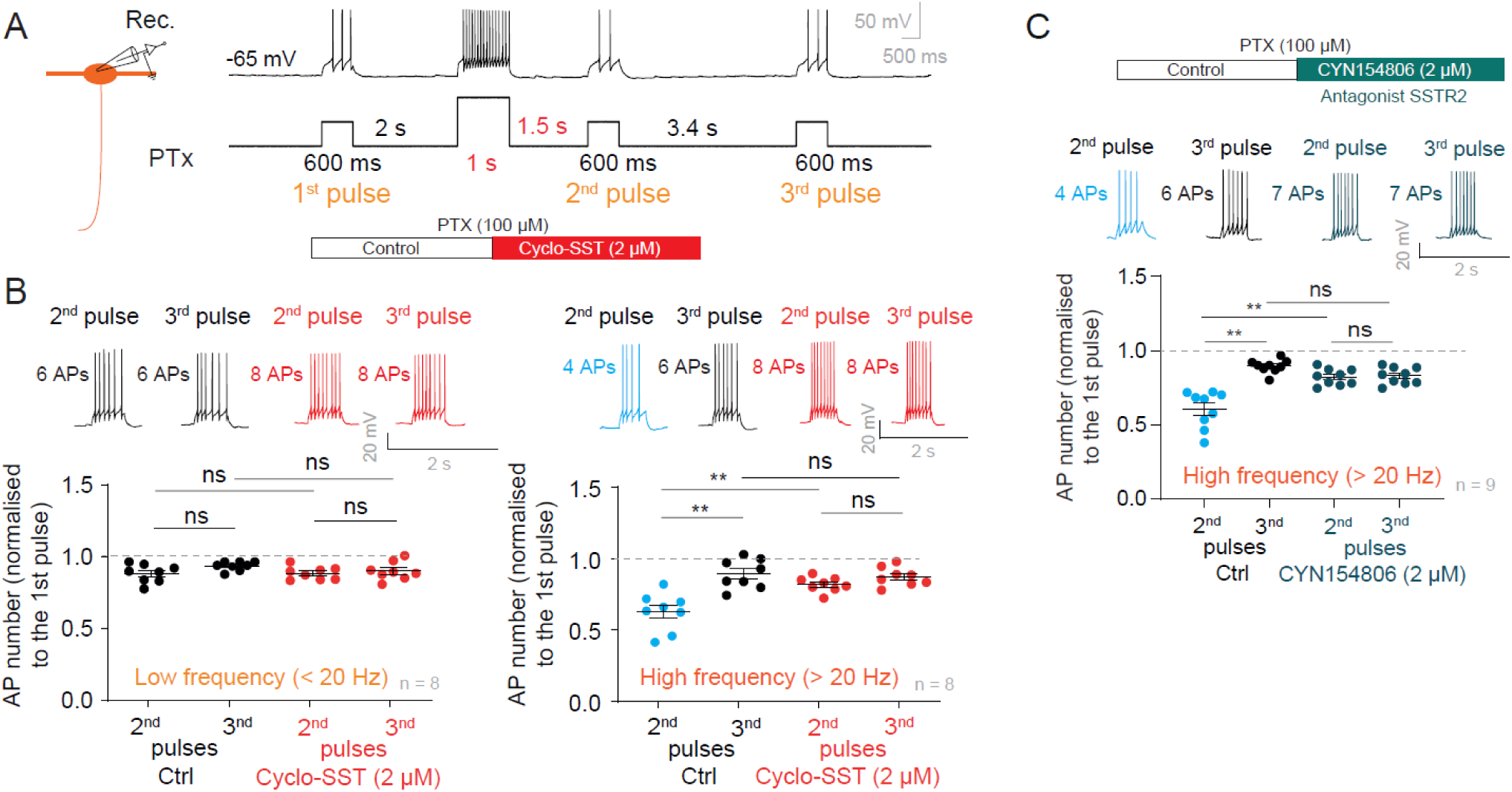
Autocrine inhibition of O-LM interneuron by SST. A. Recording configuration and stimulation protocol. B. Self-inhibition of O-LM excitability. Left, at a frequency lower than 20 Hz, the firing of the O-LM was insufficient to produce any inhibition. Right, at a frequency higher than 20 Hz, the firing frequency was inhibited (blue trace and data points). But in the presence of cyclo-SST, the inhibition was suppressed. ns, Wilcoxon test, p>0.1; **, p<0.01. C. Self inhibition of O-LM excitability induced by firing frequency larger than 20 Hz is blocked by the SST2 antagonist, CYN154806. **, Wilcoxon test, p<0.01.

We next checked whether SST release by O-LM interneurons depended on intracellular calcium. For this, BAPTA (5 mM) was added to the intracellular solution of the O-LM interneuron. In these conditions, no reduction in cell excitability was observed following O-LM firing at a frequency greater than 20 Hz (**Figure S7**), demonstrating that SST release by O-LM interneurons is calcium-dependent.

### Immunolabelling of chromogranin A and SST receptor subtypes in O-LM interneurons

As SST is thought to be released by dense core vesicles, we investigated whether SST and chromogranin A, a protein in charge of the formation of dense core vesicles, are colocalized. Immunostaining of SST and chromogranin A indicated that they were colocalized in the cell body and proximal dendrites of the O-LM cells identified based on their location and their fusiform morphology (**Figure 4A**). This strongly suggests that SST can be released by the cell body and proximal dendrites upon high-frequency firing of O-LM interneurons.

**Figure 4.**
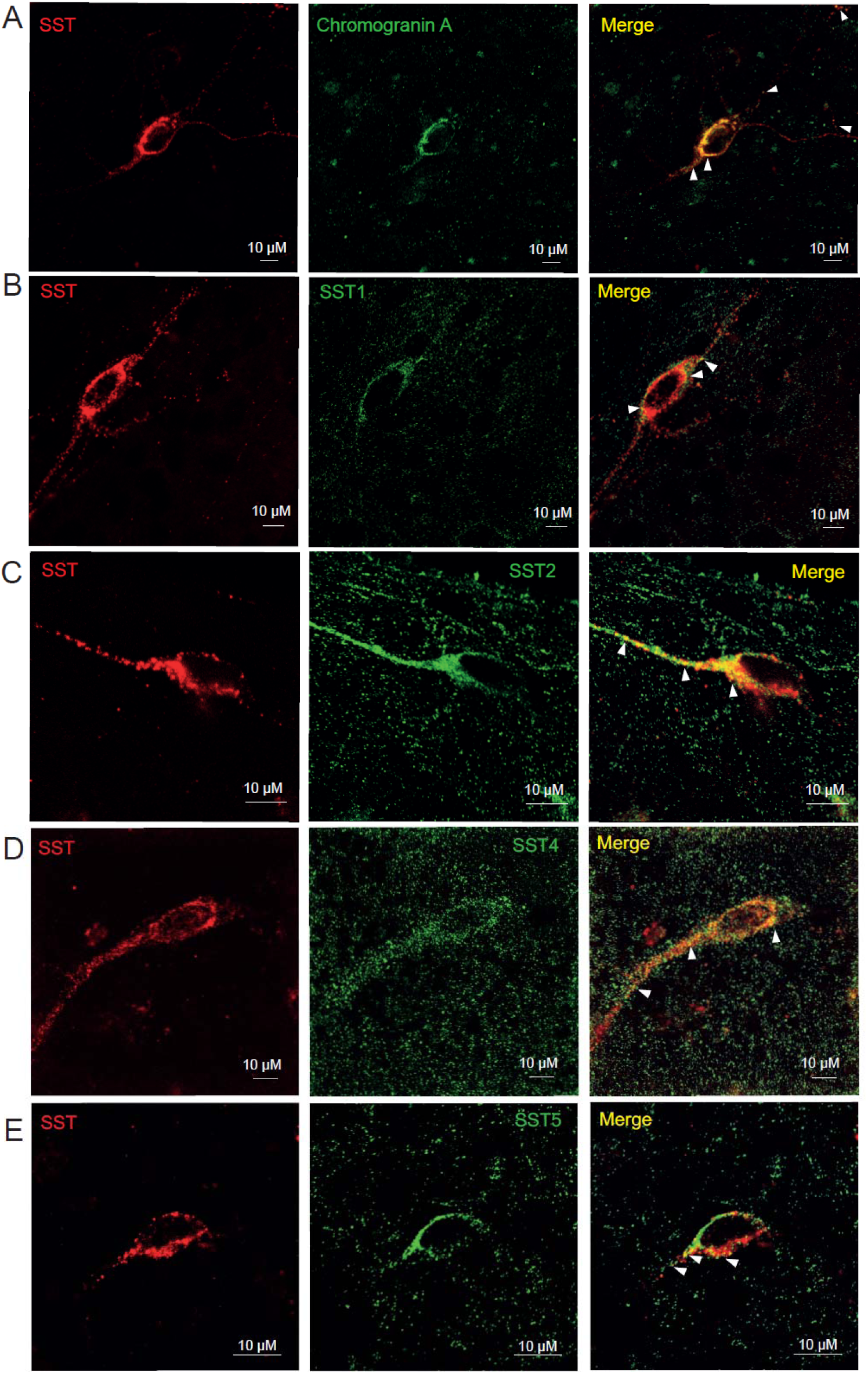
Chromagranin and SST receptors in O-LM interneurons. A. Immunostaining of Chromagranin A and SST in an O-LM cell. Left, immunostaining of SST. Middle, immunostaining of chromagranin A. Right, merge. B. Immunostaining of SST1. C. Immunostaining of SST2. D. Immunonstaining of SST4. E. Immunostatining of SST5. Arrows indicate the co-localization of SST and chromagranin (A) or SST receptors (B-E).

We next examined the subtypes of SST receptors expressed by O-LM cells using immunostaining. SST1 receptor and SST2 receptors were detected on the somas and proximal dendrites of O-LM cells, identified by their morphology and their location in the *stratum oriens* (**Figure 4B** and **4C**). SST4 immunoreactivity appeared much diffuse (**Figure 4D**) suggesting a predominantly presynaptic localization. SST5 was also present on the proximal dendrites and the cell body (**Figure 4E**). In contrast, no specific labelling was observed for SST3 (**Figure S8**) suggesting that SST3 is not expressed in the somato-dendritic compartment of O-LM interneurons. In hippocampal slice cultures, CA1 pyramidal neurons identified by phosphorylated α-CaMKII immunolabelling, expressed SST1, SST2, SST4 and SST5 receptors but not SST3 receptors on their somas and proximal dendrites (**Figure S9**). These immunostainings were performed on organotypic slice cultures of dorsal hippocampus.

### SST transiently inhibits excitatory synaptic transmission and neuronal excitability

We next tested the time-course of SST-mediated inhibition of excitatory synaptic transmission and intrinsic excitability in acutes slices of dorsal hippocampus (**Figure 5A**). Excitatory postsynaptic potentials (EPSPs) were evoked in O-LM interneurons by extracellular stimulation of CA1 pyramidal cell axons. Bath application of SST-14 transiently reduced EPSP slope during the first 2-5 minutes after which transmission recovered and exceeded the baseline level (**Figure 5B**). This reduction was associated with decrease in CV^-2^ (**Figure S10**), consistent with presynaptic mechanism. Similarly, SST-14 transiently reduced neuronal excitability during the first minutes, followed by a sustained enhancement above baseline (**Figure 5C**). The transient reduction and the sustained enhancement in intrinsic excitability were respectively due to the enhancement and reduction of the M-current (**Figure 5D**). The sustained enhancement of intrinsic excitability was also associated with a reduced rheobase (**Figure S10**).

**Figure 5.**
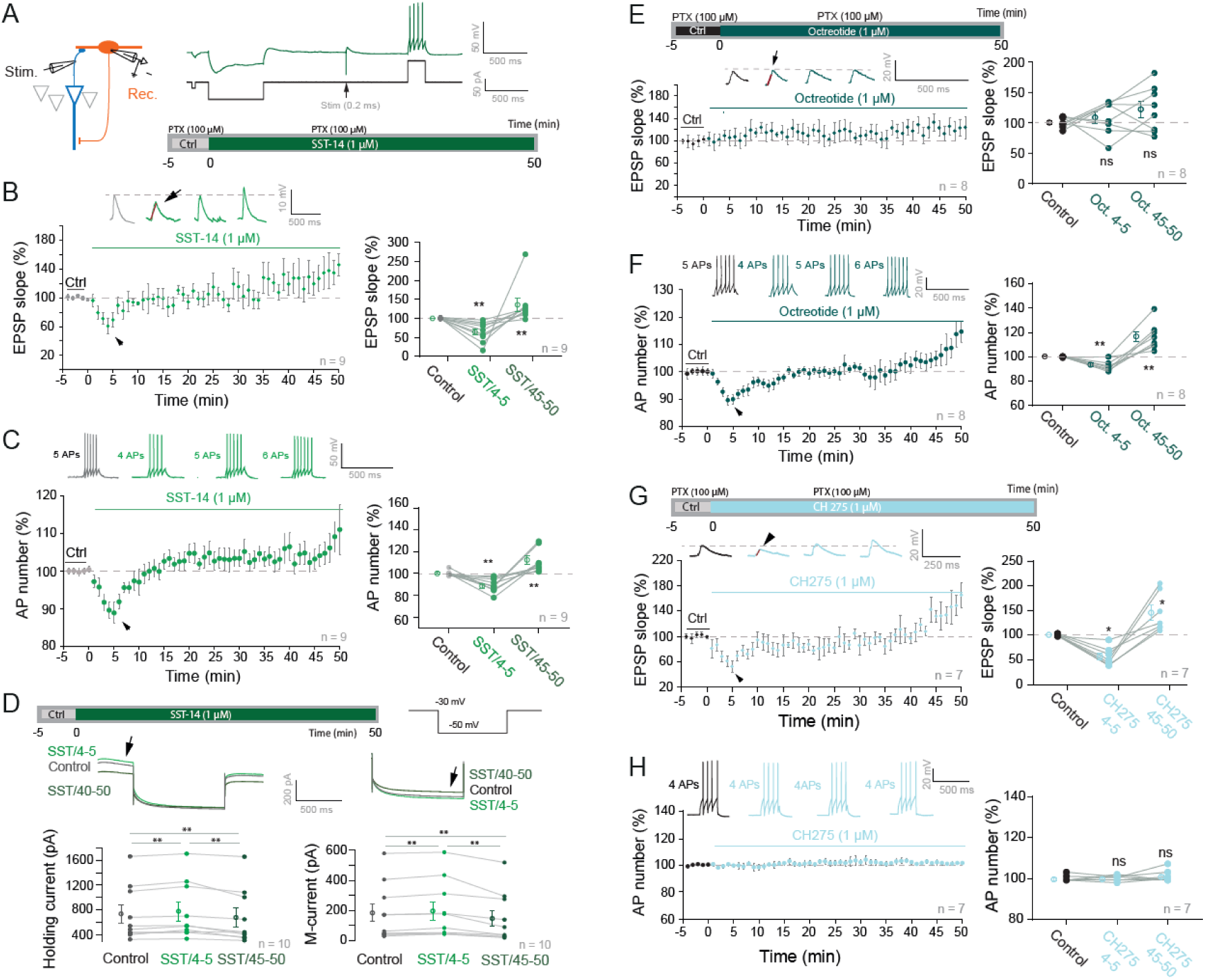
Exogeneous SST transiently inhibits both synaptic and intrinsic excitation in O-LM interneurons through SST1 and SST2. A. Experimental protocol. B. SST transiently inhibits excitatory synaptic transmission in O-LM interneurons. Left, time course showing the transient reduction in EPSP slope (black arrow) and the slow run-up. Right, plot of the transient and sustained changes. **, Wilcoxon test, p<0.01. C. SST transiently inhibits intrinsic neuronal excitability of O-LM interneurons. **, Wilcoxon test, p<0.01.D. SST enhances the Kv7-mediated M-current in O-LM interneurons. **, Wilcoxon test, p<0.01.E. In contrast to SST, the SST2 agonist octreotide (1 µM) did not reduce synaptic transmission. ns, Wilcoxon test, p>0.1. F. However, the reduction in intrinsic excitability was still present in the presence of octreotide. **, Wilcoxon test, p<0.01. G. The SST1 agonist, CH275 reduced synaptic transmission. *, Wilcoxon test, p<0.05. H. But, CH275 had no effect on intrinsic neuronal excitability. ns, Wilcoxon test, p>0.1.

The specific SST2-selective agonist, octreotide (1 µM) neither transiently inhibit nor enhanced EPSP slope at 45-50 minutes (**Figure 5E**), suggesting that the SST-dependent inhibition of excitatory synaptic transmission is mediated by another SST receptor. In contrast, octreotide reproduced the biphasic modulation of excitability observed with SST-14 (**Figure 5F**). The sustained increase in excitability induced by octreotide was also associated with a reduction of the rheobase (**Figure S11**). The SST1-selective agonist, CH275 (1 µM) transiently reduced EPSP slope by ∼40% (**Figure 5G**) without affecting postsynaptic excitability (**Figure 5H; Figure S11**), indicating that SST1 receptors are located on the presynaptic side.

We hypothesized that the transient effects of SST-14 on EPSP slope and intrinsic excitability could be mediated by the clathrin-dependent internalization of both presynaptic SST1 and postsynaptic SST2 receptors. To test this hypothesis, we preincubated slices with the clathrin-mediated endocytosis inhibitor, phenylarsine oxide (PAO, 1 µM) for 30 minutes prior to SST application (**Figure 6A**). In the presence of PAO, both synaptic transmission and intrinsic neuronal excitability remained persistently inhibited by SST-14 (**Figure 6B** and **6C**). In contrast to control conditions, the rheobase was still elevated 45-50 minutes after bath application of SST (**Figure 6D**). The time-course of SST1 and SST2 internalization was estimated by subtracting the effect of SST in PAO from the control (**Figure 6E**). We therefore conclude that SST1 and SST2 are internalized through clathrin-dependent endocytosis (**Figure 6F**).

**Figure 6.**
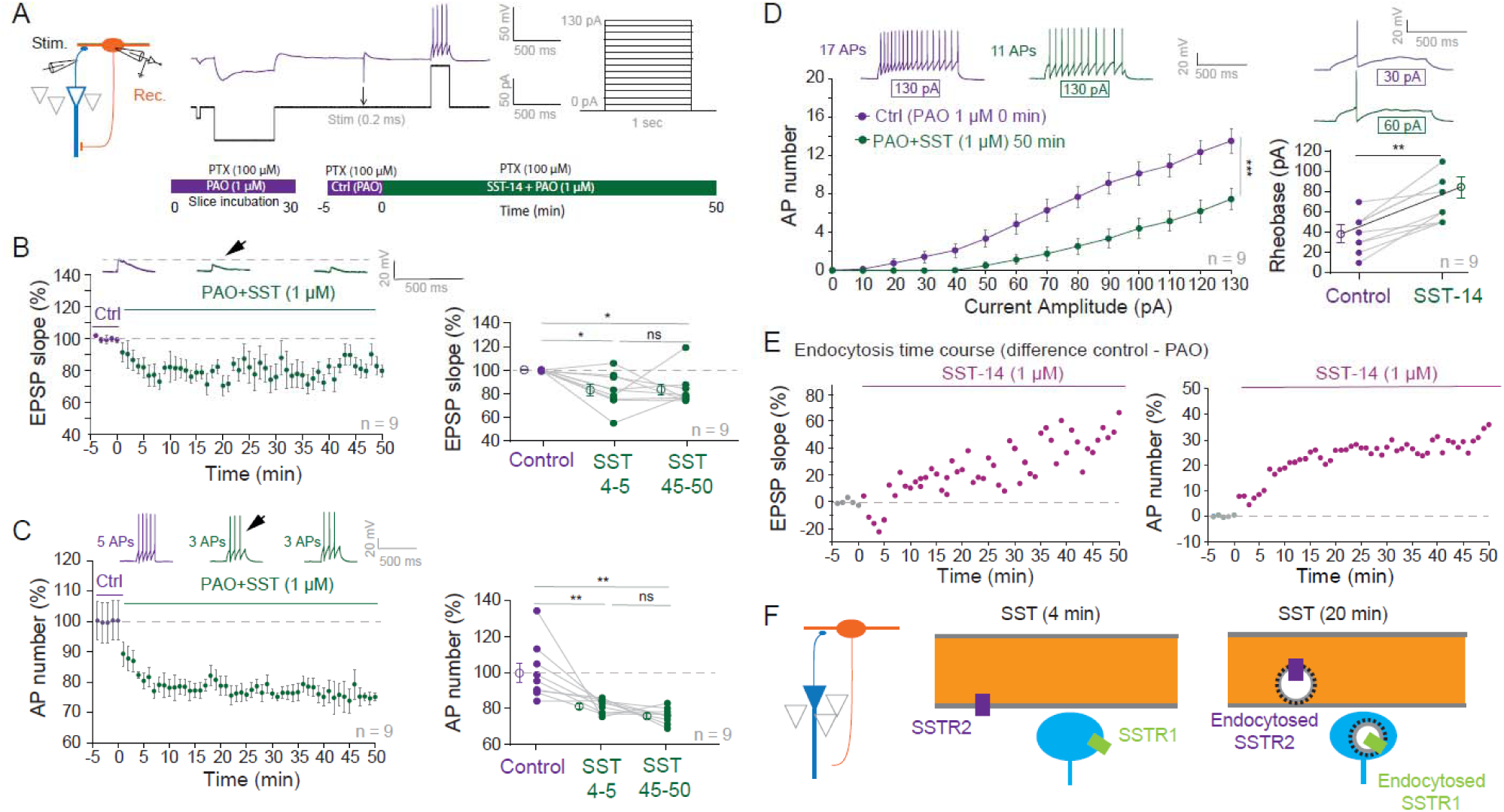
Clathrin-dependent endocytosis of SST receptors. A. Recording configuration and protocol. B. Time-course of the inhibition of excitatory synaptic transmission by SST in the presence of PAO. Note the persistent inhibition. Right, analysis of EPSP slope. ns, Wilcoxon test, p>0.05; *, p<0.05. C. Time-course of the inhibition of intrinsic excitability by SST in the presence of PAO. Right, analysis of AP number. ns, Wilcoxon test p>0.05; **, p<0.01. D. Input-output curves obtained in O-LM interneurons in the presence of the clathrin-dependent endocytosis inhibitor PAO before and after SST application. Note the reduction in intrinsic excitability and the elevation of the rheobase. **, Wilcoxon test p<0.01. E. Time-course of the effect of the SST receptor endocytosis for EPSP slope (left) and intrinsic excitability (right). F. Scheme of the endocytosis of SST receptors at the presynaptic terminal (blue) and the postsynaptic dendritic shaft (orange) at the CA1 pyramidal cell-O-LM synapse.

### Blocking SST receptors prevent LTP and LTP-IE induction

SST receptor activation enhances NMDA receptor (NMDAR) function (Pittaluga et al., 2000). We therefore tested whether blocking SST receptors with cyclo-SST reduced NMDAR-mediated EPSPs and interfere with long-term synaptic potentiation (LTP) and long-term potentiation of intrinsic excitability (LTP-IE), both of which depend on NMDAR activation in CA1 pyramidal neurons (Campanac and Debanne, 2008). NMDAR-mediated EPSPs were pharmacologically isolated with NBQX (7.5 µM). After establishing a stable baseline (**Figure S12**), cyclo-SST was also applied to the bath, resulting in a marked reduction in EPSP amplitude (51 ± 10% of the control amplitude, n = 8; **Figure 7A**).

**Figure 7.**
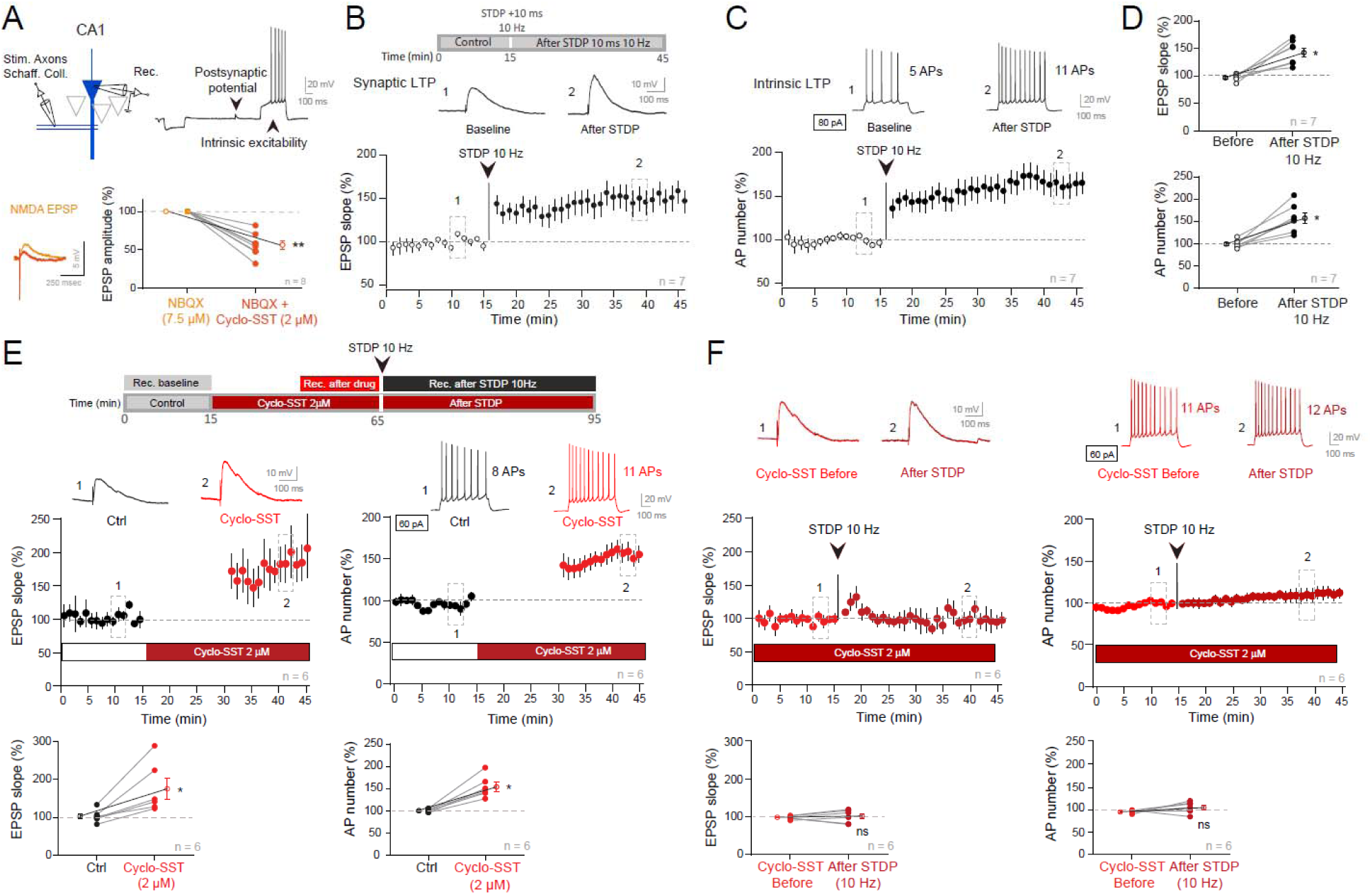
Blockade of SST receptors reduces NMDA receptor-mediated EPSP and prevents induction of both LTP and LTP-IE. A. Recording configuration and experimental protocol. **, Wilcoxon test p<0.01. B. LTP induction in control neurons by the STDP protocol. Time course of LTP and representative EPSPs before and after STDP protocol (+ 10 ms). C. LTP-IE induction in control neurons by the STDP protocol. Time course of LTP-IE and representative firing profiles before and after STDP. D. Group data of EPSP slope (top) and intrinsic excitability (bottom). *, Wilcoxon test p<0.05. E. Top, experimental protocol. Enhancement of synaptic transmission (bottom left) and intrinsic excitability (bottom right) by cyclo-SST. *, Wilcoxon test p<0.05. F. Lack of synaptic and intrinsic potentiation in the presence of cyclo-SST. ns, Wilcoxon test not significant.

LTP and LTP-IE were induced using a spike-timing-dependent plasticity (STDP) protocol pairing an evoked EPSP with a postsynaptic action potential elicited +10 ms after, at a frequency of 10 Hz (Inglebert et al., 2020). In control conditions, this protocol leads to both synaptic potentiation (144 ± 10% of the control EPSP slope, n = 7, Wilcoxon test, p < 0.05; **Figure 7B**) and intrinsic potentiation (157 ± 10% of the control spike number, n = 7; Wilcoxon test, p < 0.05; **Figure 7C** and **7D**).

Cyclo-SST application increased both baseline synaptic transmission and intrinsic excitability (respectively to 175 ± 20% and 154 ± 10%, n = 6; **Figure 7E**). However, in the presence of cyclo-SST, the STDP protocol failed to induce either LTP or LTP-IE (respectively, 102 ± 10% and 106 ± 10%, n = 6; **Figure 7F**). These results indicate that SST receptor activity is required for the induction of both synaptic and intrinsic long-lasting plasticity in CA1 pyramidal neurons. These experiments were performed on acute slices of dorsal hippocampus.

### Blocking SST receptors prevents theta oscillations

We next examined whether SST receptor activity is required for the generation of coherent theta (θ) oscillations. Robust θ oscillations were induced in intact organotypic slice cultures by bath application of 1 µM carbachol (CCh) and recorded from CA1 pyramidal neurons (**Figure 8A** and **8B**). Blocking SST receptors with cyclo-SST markedly reduced the amplitude of oscillation from 0.56 ± 0.13, n = 10 in control to 0.23 ± 0.16, n = 10 in cyclo-SST and increased the index of oscillation coherence from 0.09 ± 0.02, n = 10 to 0.21 ± 0.04, n = 10 (**Figure 8C**).

**Figure 8.**
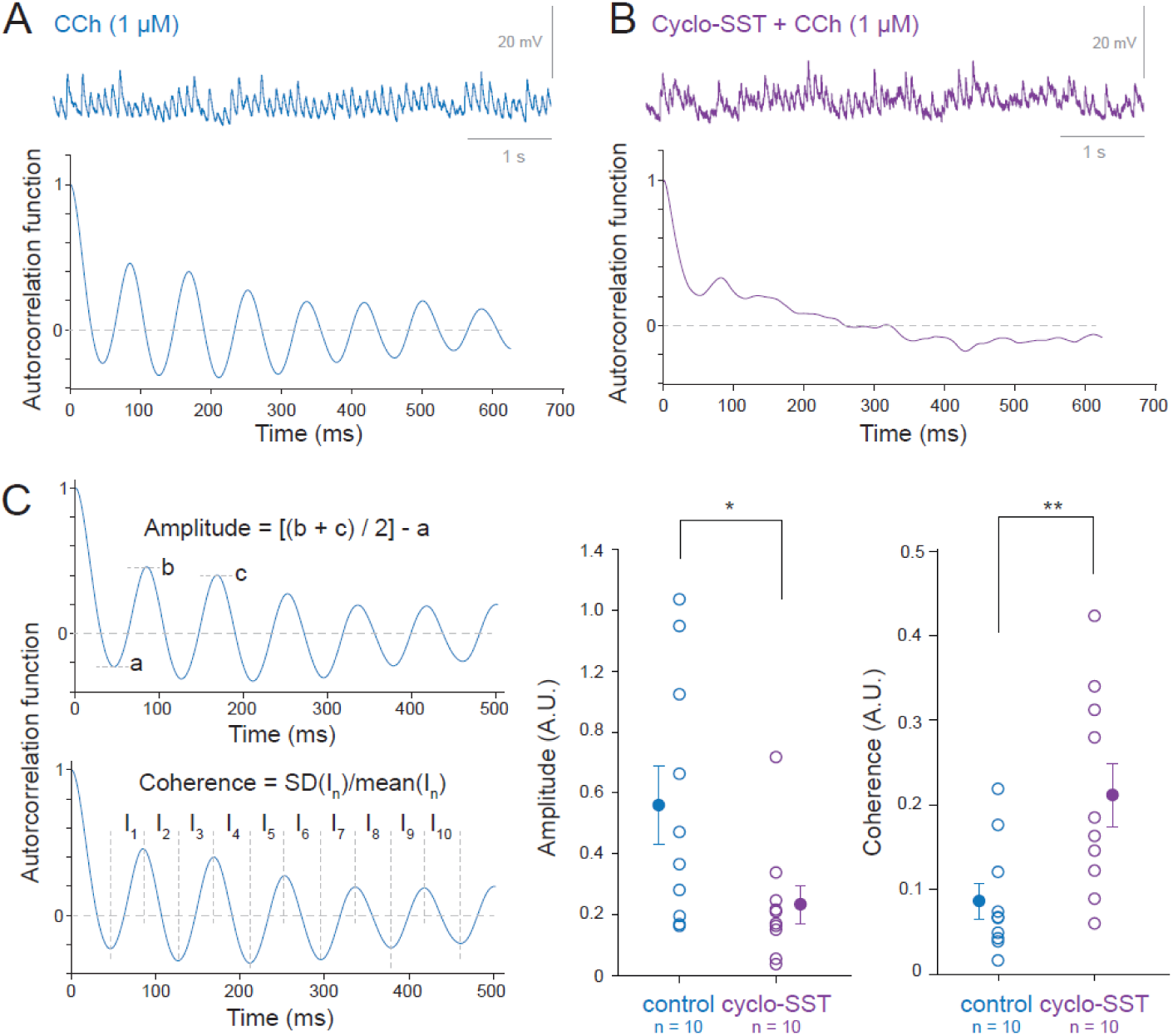
Cyclo-SST prevents oscillations at theta frequency in CA1 pyramidal neurons from organotypic slice cultures. A. Carbachol (CCh) induces oscillations of synaptic activity in the theta range. Top, representative trace. Bottom, auto-correlogram showing the persistence and the regularity of the oscillation. B. In the presence of cyclo-SST (2 µM), the synaptic activity is temporally disorganized and the auto-correlogram does not show a coherent oscillation (bottom). C. Quantitative analysis of the amplitude and coherence of the oscillation on the auto-correlogram. Left, principle of measure. Right, group data. *, Mann-Whitney U-test p<0.05; **, Mann-Whitney U-test p<0.01.

Similar effects were observed in CA1 pyramidal neurons recorded in acute slices where CCh application (25 µM) increased firing frequency in the θ frequency range under control conditions but not in the presence of cyclo-SST (**Figure S13**). These results indicate that SST receptors are necessary for the development of coherent θ-frequency oscillations in CA1 pyramidal neurons.

### Complementarity of SST- and GABA-mediated inhibition

To investigate the temporal relationship between GABA- and SST-mediated inhibition, pairs of CA1 pyramidal and O-LM cells were recorded in slice cultures of dorsal hippocampus. When an O-LM cell fires at a frequency greater than 20 Hz, the resultant postsynaptic GABAergic response in the CA1 pyramidal cell rapidly declines after ∼100 ms and reaches a stable plateau that represents nearly half of the transient response (**Figure 9A**). In contrast, SST released triggered by the same firing pattern seems to produce much slower inhibition, with significant reduction of pyramidal cell or O-LM excitability persisting ∼14 s after O-LM firing (**Figure 1B**, **Figure 2B**).

**Figure 9.**
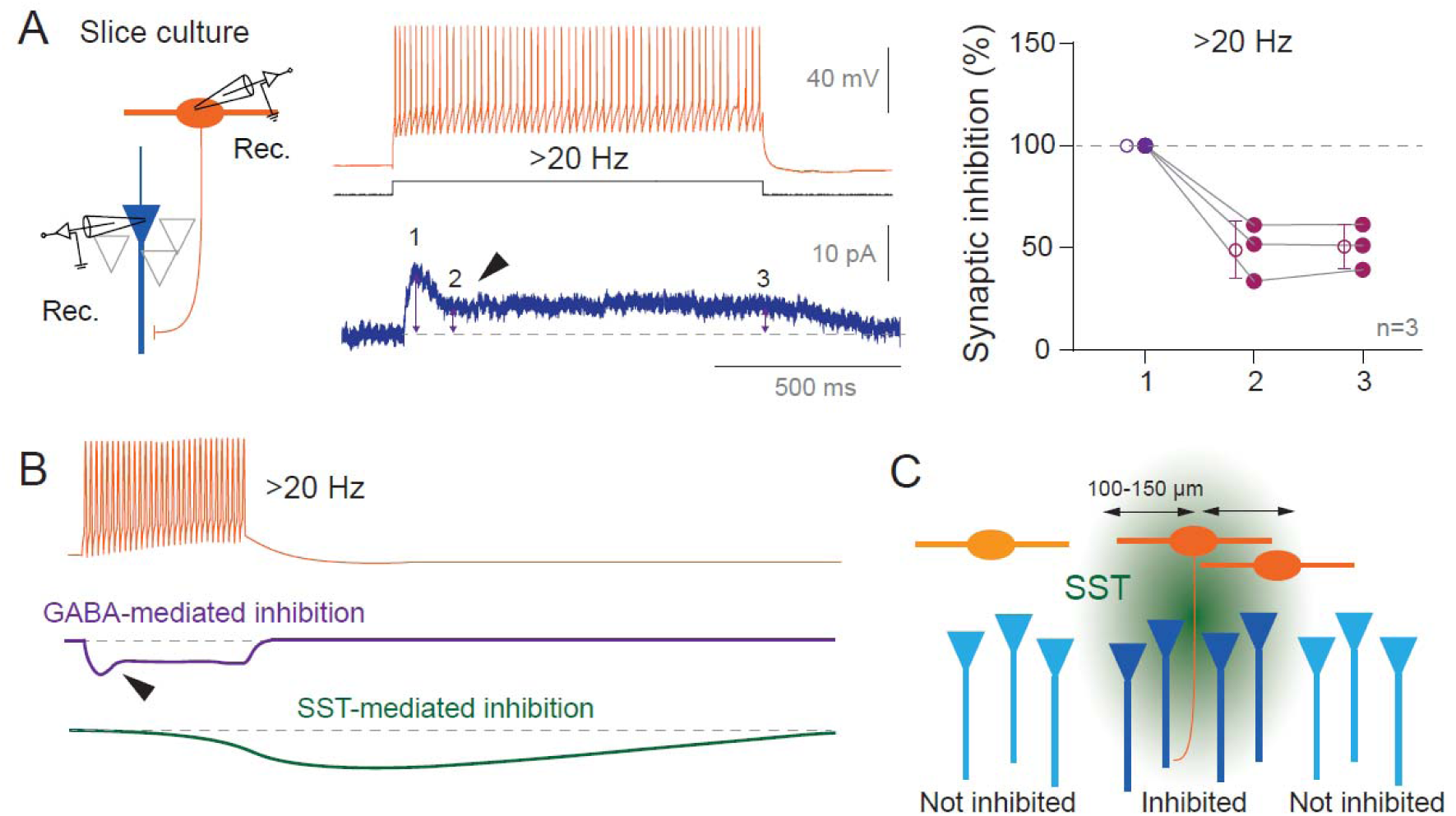
Temporal complementarity of GABA- and SST-mediated inhibitions. A. GABA-mediated inhibition attenuates overtime at O-LM / pyramidal monosynaptic connection. Left, recording configuration of an O-LM and a CA1 pyramidal neurons in organotypic slice cultures. Middle, outward GABA currents evoked by the firing at >20 Hz of a single O-LM interneuron. Right, group data at 3-point times (1: peak of the response, 2: start of the plateau, and 3: end of the plateau). B. Schematic representation of the temporal complementarity of GABA- and SST-mediated inhibitions. GABA-mediated inhibition declined during repetitive stimulation at >20 Hz whereas SST-mediated start during the train. C. Spatial extend of the inhibition produced by SST released by a single O-LM. SST inhibits pyramidal neurons, O-LM cells located near the activated O-LM and the source O-LM itself whereas it does not affect O-LM and pyramidal neurons more distant than 100-150 µm.

These findings indicate that GABA-mediated and SST-mediated inhibition appear to be complementary in time: GABA providing a fast and transient inhibition and SST, a prolonged inhibition (**Figure 9B**). SST-mediated inhibition is also spatially restricted, affecting only pyramidal neurons within a radius of ∼150 µm and O-LM cells within a radius of ∼100 µM of the release site (**Figure 9C**).

## Discussion

We show here that SST released by O-LM interneurons firing at frequency above 20 Hz inhibits the excitability of nearby CA1 pyramidal neurons and other O-LM interneurons within a spatial range of ∼100-150 µm, including the source O-LM itself through autocrine signaling. In addition, we reveal that excitatory synaptic transmission onto O-LM interneurons is also inhibited by SST. These effects are mediated by distinct receptor mechanisms: inhibition of intrinsic excitability is mediated by postsynaptic SST2 and involves M-current modulation while the inhibition of excitatory synaptic transmission onto O-LM interneurons is mediated by the presynaptic SST1 receptor. We further show that SST colocalized with chromogranin A, a protein that acts as a regulator of dense core vesicle formation and that both receptors are internalized through a clathrin-dependent process upon prolonged agonist stimulation. We also reveal that blocking SST receptors by cyclo-SST reduces NMDAR-mediated responses, induces both synaptic and intrinsic potentiation in CA1 pyramidal cells and abolishes the induction of further synaptic and intrinsic potentiation. In addition, SST receptor blockade prevents the development of theta oscillations. Collectively, these results indicate that SST function is critical for maintaining normal hippocampal function across multiple temporal and spatial scales.

### Requirements for SST release by single O-LM interneurons

O-LM interneurons firing at frequencies above 20 Hz were able to inhibit both CA1 pyramidal neurons and other neighboring O-LM cells. Such firing frequency is commonly observed in O-LM cells during network activity *in vivo* (Klausberger et al., 2003; Varga et al., 2012) and *in vitro* (the present study). Neuropeptide release from dense core vesicles classically depends on the rate of stimulation and requires repetitive stimulations (Dutton and Dyball, 1979; Whim and Moss, 2001). Thus, SST released by single interneurons follows the same principle.

SST-mediated inhibition of intrinsic excitability occurs within a radius of ∼150 µm for pyramidal neurons and ∼100 µm for O-LM interneurons. Interestingly, the inhibition for both cell types followed a linear relationship with the distance. SST2 receptors which mediate this inhibition are located on the cell bodies and the dendrites of CA1 pyramidal cells and O-LM interneurons, ensuring that SST released by a single O-LM cell can act on both neuronal types. Whether other interneuron classes such as parvalbumin (PV)-positive basket cells, cholecystokinin interneurons, vasointestinal peptide-containing (VIP) interneurons or bistratified interneurons are similarly inhibited by SST remains unknown. However, this is possible as neocortical interneurons such as PV or VIP interneurons express high levels of SST receptor transcripts (Tasic et al., 2018; Song et al., 2021).

### Time course of SST-mediated inhibition

The precise onset of SST-mediated intrinsic inhibition is difficult to assess, but our data indicate that it begins within 2.5 seconds after the end of stimulation and persists at least 14 s after in both CA1 pyramidal neurons and O-LM interneurons. So, the time-course of SST inhibition is considerably slower than that of any classical inhibitory neurotransmitter. Once released, SST is degraded within 1-3 min. The relatively slow action of SST may result from the prolonged fusion kinetics of dense-core vesicles (Kreutzberger et al., 2019) and the metabotropic activation of SST2 to inhibit postsynaptic Kv7 channels by lipid synthesis (Incontro et al., 2025).

Autocrine SST-mediated inhibition of O-LM interneurons exhibits faster kinetics, with the maximal reduction observed at 1.5 seconds after the conditional train of action potentials and return to near-control levels by 4.9 seconds. The reason for this faster time-course in autocrine inhibition has not been investigated here. Nevertheless, the time required for an SST molecule to reach a proximal target (here, its own dendrite) must be shorter than a distant target (i.e., a different cell) according to Brownian dynamics. Additionally, SST2 internalization might proceed more rapidly during autocrine signaling, contributing to the faster decay of inhibition.

### SST receptors in O-LM interneurons

We identified four SST receptor subtypes - SST1, SST2, SST4 and SST5 - in O-LM interneurons. SST1 displayed minimal co-localization with SST within O-LM cells, consistent with a predominant presynaptic location. Supporting this hypothesis, selective activation of SST1 with CH275 reduced synaptic excitatory synaptic transmission without affecting intrinsic excitability. SST1 undergoes clathrin-dependent endocytosis as indicated by the persistence of the inhibition of excitatory synaptic transmission in the presence of a clathrin-endocytosis inhibitor.

SST2 was found to be co-localized with SST in O-LM cells, suggesting its postsynaptic location. Supporting this hypothesis, we found that specifically stimulating SST2 with octreotide reduced postsynaptic intrinsic excitability but not synaptic transmission. Like SST1, SST2 also underwent endocytosis upon sustained activation, as the inhibition of clathrin-dependent endocytosis revealed a persistent inhibition of intrinsic excitability. While all SST receptors except SST4 can undergo endocytosis in dissociated cells (Csaba and Dournaud, 2023), our data provide, to our knowledge, the first evidence for the internalization of SST1 in native tissue.

SST4 was detected at the periphery of O-LM interneurons, indicating a presynaptic location, whereas SST5 was colocalized with SST, suggesting a postsynaptic expression. In contrast, no labeling of SST3 has been observed. The role of SST4 and SST5 in the physiology of O-LM interneurons remain unexplored here due to the lack of available and selective antagonists of these receptors.

Constant stimulation of the SST2 receptor by SST or octreotide causes its internalization through clathrin-dependent endocytosis (Csaba et al., 2001; Lelouvier et al., 2008). This SST2 endocytosis elevates intrinsic excitability in O-LM cells. However, the blockade of clathrin-dependent endocytosis with PAO prevented this increased excitability.

### SST receptor blockade prevents induction of LTP and LTP-IE in pyramidal cells

We show that receptor blockade by cyclo-SST enhances both synaptic transmission and intrinsic excitability in CA1 pyramidal neurons, while preventing further induction of synaptic and intrinsic plasticity. The increase in synaptic transmission and intrinsic excitability is likely due to the removal of the tonic inhibition exerted by SST on synaptic transmission and intrinsic excitability. Because the NMDA receptor activation is a critical step for induction of both STDP (Gustafsson et al., 1987; Debanne et al., 1994; Bi and Poo, 1998) and intrinsic plasticity associated with LTP (Campanac and Debanne, 2008), the known ability of SST to potentiate NMDA receptor function (Pittaluga et al., 2000, 2021) suggests mechanistic link. Thus, blockade of SST receptors relieves the tonic up-regulation of the NMDA receptor function and impairs further induction of LTP and LTP-IE in pyramidal neurons.

Intrinsic excitability in O-LM interneurons is bidirectionally tuned in parallel with long-lasting synaptic plasticity (Incontro et al., 2021; Sammari et al., 2022). It follows that release of SST would be persistently facilitated after LTP-IE induction while LTD-IE may be associated with reduced SST.

### Inhibition of ***θ*** oscillations by cyclo-SST

θ oscillations were deeply altered by blocking SST receptors with cyclo-SST. In fact, θ oscillations depend on a delicate interplay between intrinsic neuronal properties and multiple synaptic interactions established between interneurons and pyramidal neurons (Bezaire et al., 2016). Disruption of this balance such as by h-channel blockers prevents θ oscillations in the hippocampus and the neocortex (Gastrein et al., 2011). Synaptic stimulation at θ frequency induces synaptic plasticity and intrinsic plasticity in pyramidal cells (Mulkey and Malenka, 1992; Brager and Johnston, 2007; Gasselin et al., 2017) and in O-LM interneurons (Sammari et al., 2022).

Following cyclo-SST application, the persistent elevation in excitability and synaptic transmission observed in both pyramidal cells and O-LM interneurons, combined with the loss of plasticity in pyramidal neurons could contribute in part to the observed disruption of θ oscillations. The cyclo-SST-induced reduction of the NMDAR component may further exacerbate this effect, since NMDAR activation is known to be also critical for θ rhythm generation in the hippocampus (Kazmierska and Konopacki, 2013; Gu et al., 2017).

### Generalization of SST release by interneurons

SST is present in several subtypes of hippocampal interneurons, including O-LM interneurons, bistratified cells and oriens-oriens interneurons (Klausberger and Somogyi, 2008; Müller and Remy, 2014; Chamberland et al., 2024) as well as in neocortical interneurons such as Martinotti and non-Martinotti cells (Park et al., 2025). The principles described here may therefore extend beyond O-LM cells, although further experiments are needed to confirm this.

## Conclusion

Our findings identify SST release from single O-LM interneurons as a potent modulator of hippocampal excitability and circuit dynamics within a restricted spatial area and persisting for several seconds, thus distinguishing SST signaling from classical fast neuro-inhibition. By tonically regulating NMDAR function, SST maintains the capacity for synaptic and intrinsic plasticity and supports the balance of excitation and inhibition necessary for θ oscillations.

Together, these results establish SST as a gatekeeper of CA1 circuit processing, with major implications for learning, memory and rhythm generation. Dysregulation of SST signaling – whether through altered release, receptor function, or endocytosis – may therefore contribute to network dysfunction in neurological and psychiatric disorders.

## Material and methods

### Acute slices of rat hippocampus

All experimental procedures followed institutional guidelines for the care and use of laboratory animals (Council Directive 86/609/EEC and French National Research Council) and were approved by the local health authority (Préfecture des Bouches-du-Rhône, Marseille). Briefly, 14-to 21-day-old Wistar rats (Charles River) of either sex were anesthetized with isoflurane and decapitated. Hippocampal slices (350 µm) from the dorsal hippocampus were cut using a parasagittal orientation in a N-methyl-D-glucamine (NMDG)-solution containing (in mM): 92 NMDG, 2.5 KCl, 1.2 NaH_2_PO_4_, 30 NaHCO_3_, 20 HEPES, 25 Glucose, 5 Sodium Ascorbate, 2 Thiourea, 3 Sodium Pyruvate, 10 MgCl_2_, 0.5 CaCl_2_ on a vibratome (Leica VT1200S). They were transferred in an NMDG-solution for 10 min maintained at 37°C before resting for 1h at room temperature in oxygenated (95% O_2_/5% CO_2_) artificial cerebro-spinal fluid, ACSF (in mM: 125 NaCl, 2.5 KCl, 0.8 NaH_2_PO_4_, 26 NaHCO_3_, 3 CaCl_2_, 2 MgCl_2_ et 10 Glucose).

### Organotypic slice cultures of hippocampus

Slices cultures were prepared as described previously (Debanne et al., 2008; Extrémet et al., 2023). Young Wistar rats (P6–P8) were killed by decapitation, the brain was removed, and each hippocampus dissected. Hippocampal slices (350 μm) were obtained using a Vibratome (Leica, VT1200S). They were placed on 20-mm latex membranes (Millicell) inserted into 35-mm Petri dishes containing 1 ml of culture medium and maintained for up to 10-20 days in an incubator at 35.5 °C, 95% O_2_–5% CO_2_. The culture medium contained (in ml) 25 MEM, 1.25 HBSS, 12.5 horse serum, 0.5 penicillin/streptomycin, 0.8 glucose (1 M), 0.1 ascorbic acid (1 mg/ml), 0.4 Hepes (1 M), 0.5 B27, and 8.95 sterile H_2_O. Slice cultures were used i) to assess the presence of dense core vesicles and the five SST receptors in O-LM interneurons using immunohistochemistry, ii) to test the role of cyclo-SST on carbachol-induced oscillations at θ frequency and iii) to record pairs of synaptically connected pyramidal and O-LM neurons.

### Electrophysiology and data acquisition

Because synaptic inhibition was blocked by picrotoxin (PiTX, 100 µM), the CA3 area was removed surgically to avoid epileptiform activity for all the experiments except when carbachol-induced oscillations at θ frequency were recorded which requires an intact hippocampal circuit. Each slice was transferred to a temperature-controlled (30°C) recording chamber with oxygenated ACSF + PiTx. O-LM hippocampal interneurons were identified by the location of their soma (*stratum oriens* of the CA1), their morphology (a spindle-shaped cell body horizontally oriented along the pyramidal layer) and their peculiar electrophysiological signature (the characteristic “sag” depolarizing potential in response to hyperpolarization currents injections and the typical “saw-tooth” shape of action-potential after-hyperpolarization). Whole-cell patch-clamp recordings were obtained from CA1 O-LM interneurons with electrodes filled with an internal solution containing (in mM): K-gluconate, 120; KCl, 20; HEPES, 10; EGTA, 0; MgCl_2_6H_2_O, 2; Na_2_ATP, 2; spermine, 0.1. In the case of dual recording from a pyramidal cell and an O-LM interneuron or from 2 O-LM interneurons, neuron types were identified on the base of their firing profile and their morphology. Stimulating pipettes filled with extracellular saline solution were placed in the stratum oriens to stimulate the axons of CA1 pyramidal cells.

Recordings were obtained using a Multiclamp 700B (Molecular Devices) amplifier and pClamp10.4 software. Data were sampled at 10 kHz, filtered at 3 kHz, and digitized by a Digidata 1440A (Molecular Devices). All data analyses were performed with custom written software in Igor Pro 6 (Wavemetrics). Values of membrane potential were not corrected for liquid junction potential (∼ -12 mV).

Apparent input resistance (R_in_) was tested by current injection (−120 pA; 800 ms); EPSPs were evoked at 0.1 Hz and the stimulus intensity (100 µs, 40–100 µA) was adjusted to evoke subthreshold EPSPs (4–10 mV). Synaptic transmission was measured between 2 and 4 ms from the onset of the EPSP. Short current step injections (+70/+120 pA, 250 ms) were applied at each sweep in order to measure the intrinsic excitability as the number of spikes over time. Series resistance was monitored throughout the recording and only experiments with stable resistance were kept (changes <10%). Before and after SST application in the presence of either PAO, cyclo-SST, CYN, octreotide, SST or CH275, a protocol was designed in order to plot input-output curves by measuring AP number in response to incrementing steps of current pulses.

Simultaneous paired recordings from O-LM and CA1 pyramidal neurons or from two O-LM interneurons were obtained in acute slices. In addition, paired recordings from connected O-LM to CA1 pyramidal neurons were obtained in organotypic hippocampal slice cultures.

### Induction of theta oscillations

Theta oscillations were induced in acute slices of the dorsal hippocampus and in hippocampal slice cultures with bath application of CCh (25 µM and 1 µM respectively). Oscillations appeared 2-5 minutes after CCh application.

### Analysis of subthreshold oscillations

Autocorrelation function was obtained from the oscillating membrane potential in ClampFit. Amplitude and temporal coherence of oscillations were analyzed as previously established (Bringuier et al., 1997; Gastrein et al., 2011). Briefly, the optima in the autocorrelogram were detected and the amplitude of oscillation was determined by the amplitude between the first minimum and the average of the first two maxima. The coefficient of temporal oscillation coherence was determined as the ratio of the standard deviation over 10 intervals divided by the mean.

### Pharmacology

Excepted BAPTA (1,2-Bis(2-aminophenoxy) ethane-N,N,N’,N’-tetraacetic acid tetrakis(acetoxy-methyl ester)) that was added to intracellular solution, all drugs were bath applied. Picrotoxin was purchased from Abcam. BAPTA, SST-14, Cyclosomatostatin, CYN154806, Octreotide, NBQX, Carbachol (Carbamoylcholine chloride) were obtained from Tocris. Phenylarsine oxide (PAO) was obtained from Sigma-Aldrich and CH275 from MedChemExpress.

### Voltage-clamp measurements Kv7 channel-mediated current

To measure the M-current mediated by Kv7 channels in the absence and presence of SST in O-LM interneurons, voltage steps from -30 mV to -50 mV were applied (Lawrence et al., 2006), and the relaxing component of the M-current was then measured before and after SST.

### Morphological identification of biocytin-filled O-LM interneurons

O-LM interneurons and CA1 pyramidal neurons were filled with biocytin to visualize the dendritic and axonal morphology of the recorded neurons. Biocytin (0.2-0.4%, Sigma Aldrich) was added to the pipette solution, and cells were filled at least for 20 min. Biocytin was revealed with streptavidin complex coupled to Alexa Fluor 594 (Thermo Fisher Scientific), and examined using confocal microscopy (Zeiss, LSM 780). The morphology of the neurons was reconstructed using ImageJ.

### Induction of LTP and LTP-IE

LTP and LTP-IE were induced using a +10 ms pairing between the evoked EPSP and the postsynaptic action potential that was repeated 100 times at 10 Hz (Campanac and Debanne, 2008; Inglebert et al., 2020). The slope of the EPSP and the AP number were then analyzed before and after the pairing protocol and normalized on the control period.

### Immunofluorescence of organotypic hippocampal slice cultures

To perform immunostainings, 10 DIV organotypic slices were rinsed in PBS at 37°C and immediately immersed in ice-cold 4% paraformaldehyde in PBS. Slices were fixed at 4°C for 15 minutes and subsequently incubated in PBS containing 50 mM NH4Cl (15 min, RT) to quench the remaining free aldehyde groups. Afterwards, the Millicell membrane containing the slices was cut with a scalpel, and the slices were blocked and permeabilized (O/N, 4°C) in PBS containing 0.5% Triton X-100 and 5% Normal Goat Serum (NGS, Thermo Fisher Scientific). Then, slices were incubated with the primary antibodies (24h, 4°C) in a solution containing 0.5% Triton X-100 and 2% NGS in PBS. The following primary antibodies were used in these experiments: mouse anti-SST (1:500, Thermo Fisher Scientific 14-9751-82), mouse anti-Phosph-CaMKII alpha (Thr286; 1:100, Thermo Fisher Scientific MA1-047), rabbit anti-chromogranin A (1:200, Synaptic Systems 259 003), rabbit anti-SST1 (1:500, Thermo Fisher Scientific, PA3-206), rabbit anti-SST2, SST3, SST4 and SST5 (1:100, 1:200, 1: 400, 1:200, Alomone Labs, ASR-006, ASR-003, ASR-004 and ASR-005). Afterwards, sections were washed three times (20 min each) in PBS containing 0.5% Triton X-100 and then incubated with the appropriate secondary antibodies (2 h, RT) in PBS containing 0.5% Triton X-100 and 2% NGS. The following secondary antibodies were used in these experiments: goat anti-rabbit Alexa Fluor 488 (1:600, Jackson ImmunoResearch), goat anti-mouse Alexa Fluor 594 (1:500, Jackson ImmunoResearch). Finally, sections were washed three times (20 min each) in a PBS solution containing 0.5% Triton X-100, then incubated with DAPI (1.5 µg/mL in PBS, Sigma-Aldrich) for 10 min, washed one last time in PBS and mounted in Vectashield Antifade Mounting Medium (Vector laboratories). Slides were kept at 4°C until use. The double immunostaining using anti-SST and either anti-SST2, anti-SST4 or anti-SST5 antibodies were performed using the abovementioned steps but without permeabilizing agent to obtain a clearer extracellular staining.

### Immunofluorescence image acquisition and analysis

Immunofluorescence images were acquired on a Zeiss LSM-780 confocal scanning microscope. All experiments images were analysed with ImageJ.

### Statistical analysis

Pooled data are presented as mean ± SEM. Statistical comparisons were made using Wilcoxon or Mann-Whitney test as appropriate with Prism (GraphPad) software. Differences were considered as significant when p < 0.05 (*p < 0.05; **p < 0.01; ***p < 0.001).

## Acknowledgments

We thank K. Milton and A. Venture for excellent animal care and N Boumedine-Guignon for excellent assistance.

## Author contributions

M.L.M. designed and performed research, analyzed the data, built the figures and wrote the paper. M.S. performed research and analyzed the data. C.A. performed research and analyzed the data. A.L.C. performed research and analyzed the data. M.R. performed research and analyzed the data. D.D. designed research, analyzed the data, provided funding, built the figures and wrote the paper. S.I. designed and performed research, analyzed the data and edited the manuscript.

## Data availability

All data needed to evaluate the paper are present in the manuscript and/or in the Supplementary Materials.

---

**Figure S1.**
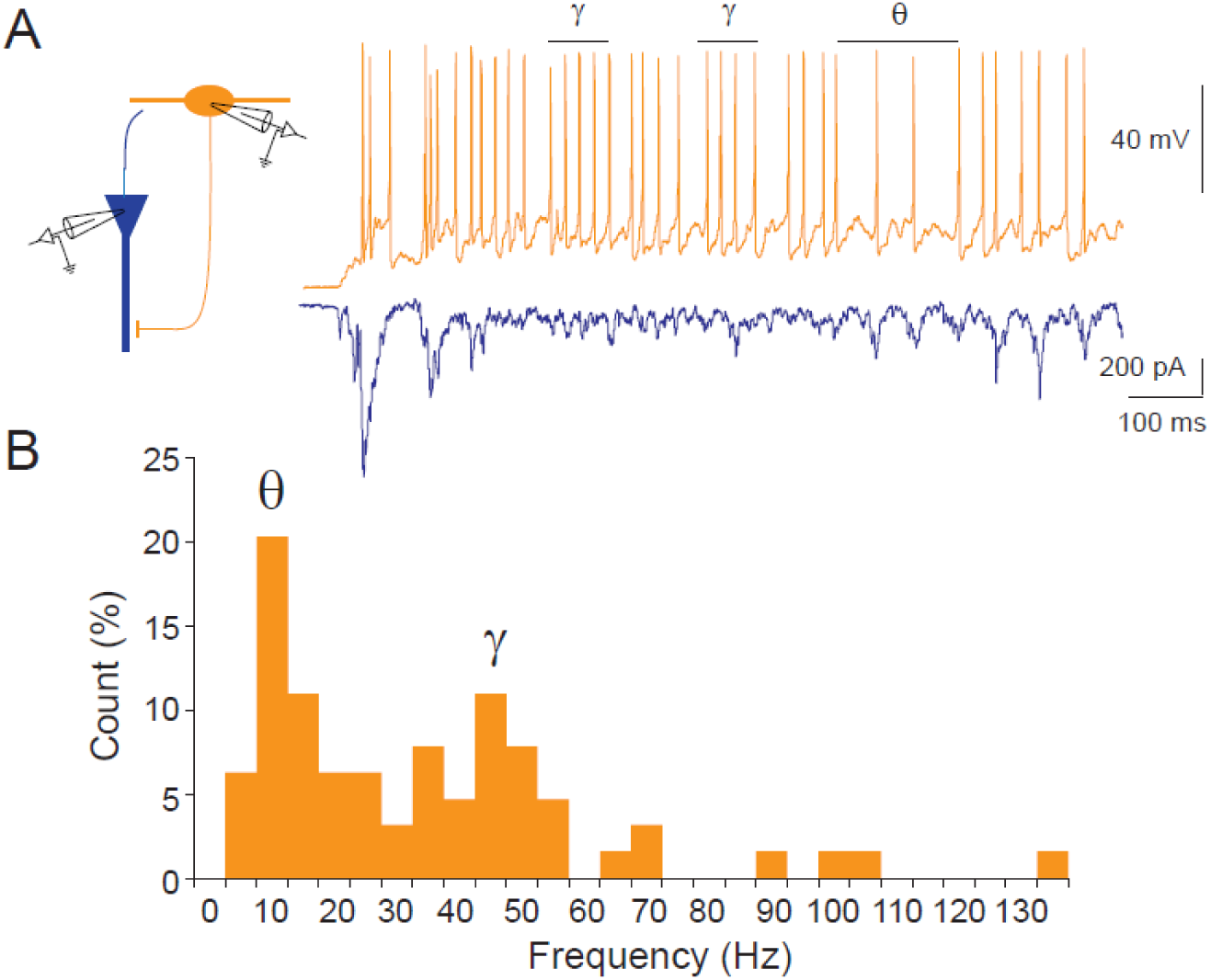
Firing of an O-LM interneuron above 20 Hz during spontaneous network activity and frequency-dependent inhibition of CA1 pyramidal cell firing. A. Recording configuration in an organotypic slice culture (left) and representative traces (right) from an O-LM cell (orange trace in current clamp) and a CA1 pyramidal neuron (blue trace in voltage-clamp) recorded simultaneously. The O-LM interneurons fires during ∼1 s at a mean frequency above 20 Hz. B. Frequency histogram of spiking activity measured in the O-LM interneuron. Activity in the γ frequency range represents about half of the total spiking activity.

**Figure S2.**
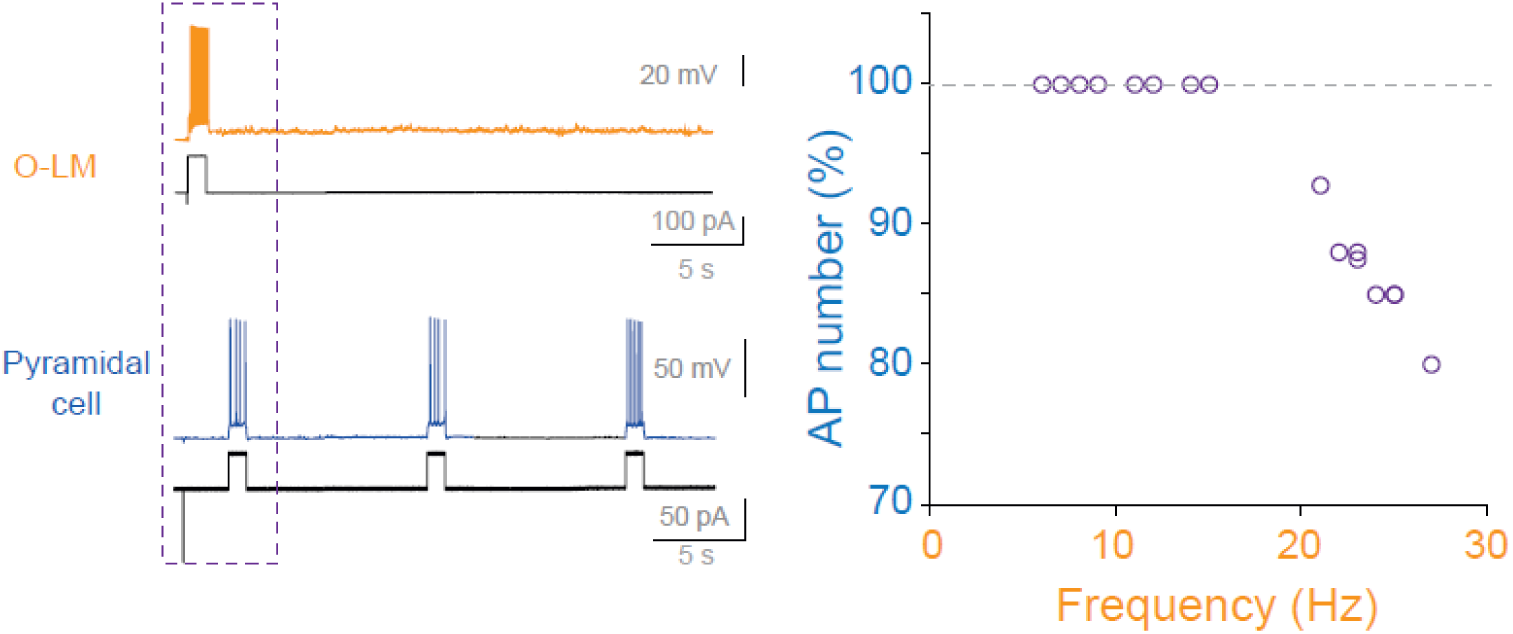
**CA1 pyramidal cell firing inhibition depends on firing frequency in the O-LM cell. Left, protocol of SST released by the O-LM cell (orange) on the excitability of the pyramidal cell (blue). Right, AP number in the pyramidal cell as a function of O-LM firing frequency.**

**Figure S3.**
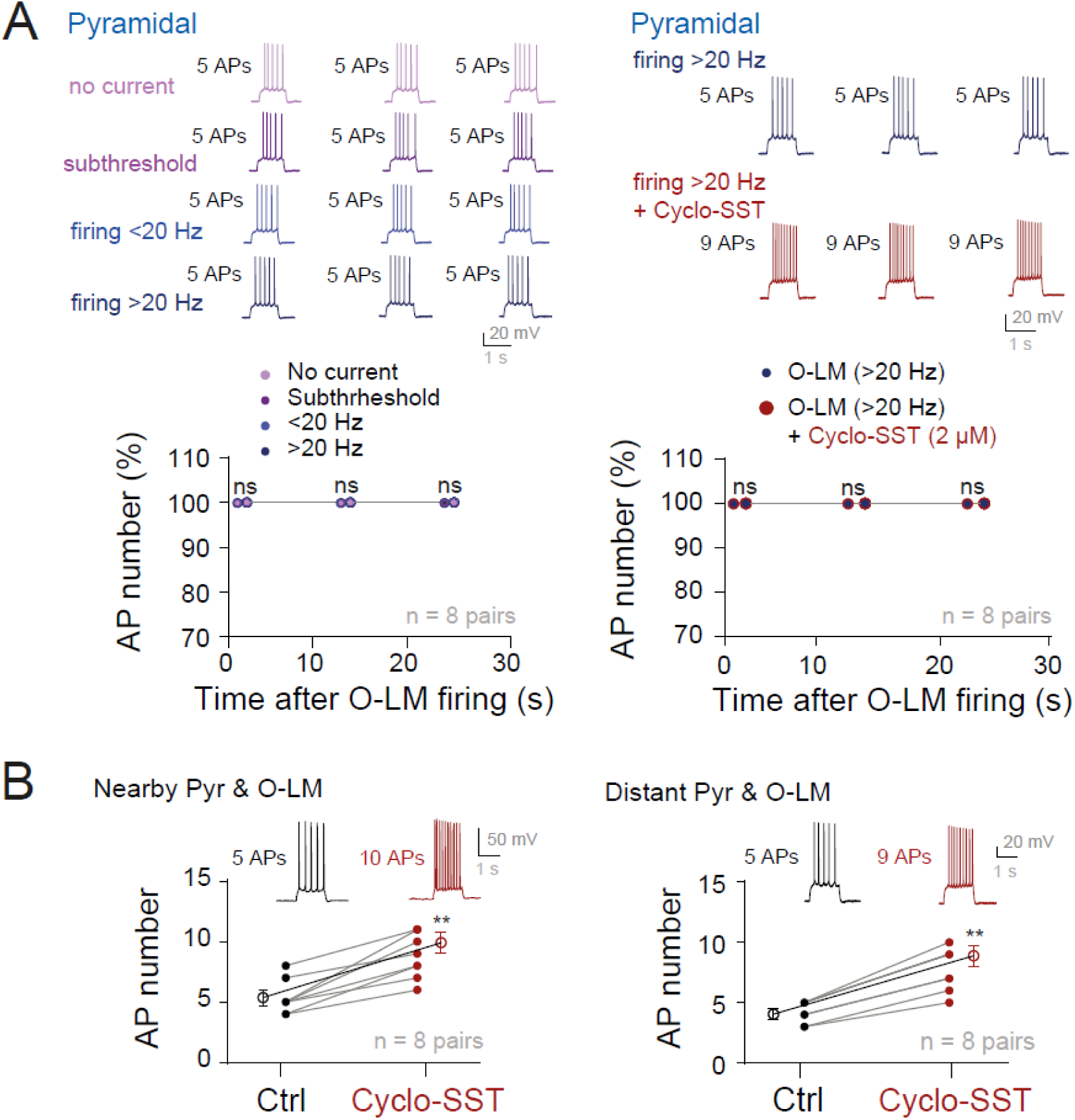
Lack of inhibition at distant pyramidal/O-LM cells and enhanced excitability induced by cyclo-SST in pyramidal neurons. A. Left, firing in the O-LM did not inhibit CA1 pyramidal cell firing when the two neurons were distant. Right, same experiment in the presence of cyclo-SST (2 µM). ns, Wilcoxon test not significant. B. Elevated firing in pyramidal neurons induced by cyclo-SST when the pyramidal cell and the O-LM interneuron were nearby (left) or distant (right). **, Wilcoxon test p<0.01.

**Figure S4.**
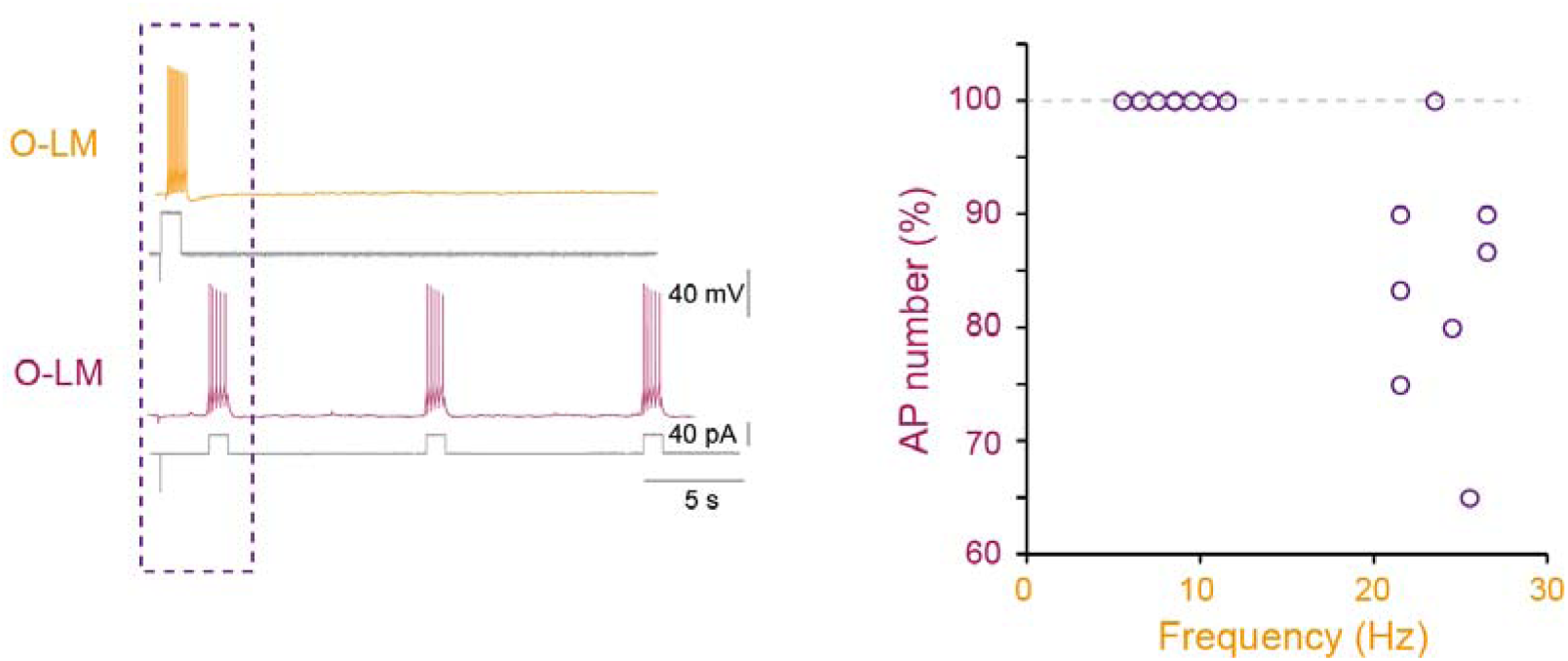
Inhibition of O-LM firing depends on firing frequency in another adjacent O-LM cell. Left, protocol of SST released by the first O-LM cell (orange) on the excitability of the second O-LM (purple). Right, AP number in the second O-LM cell (purple) as a function of O-LM firing frequency in the other adjacent O-LM (orange).

**Figure S5.**
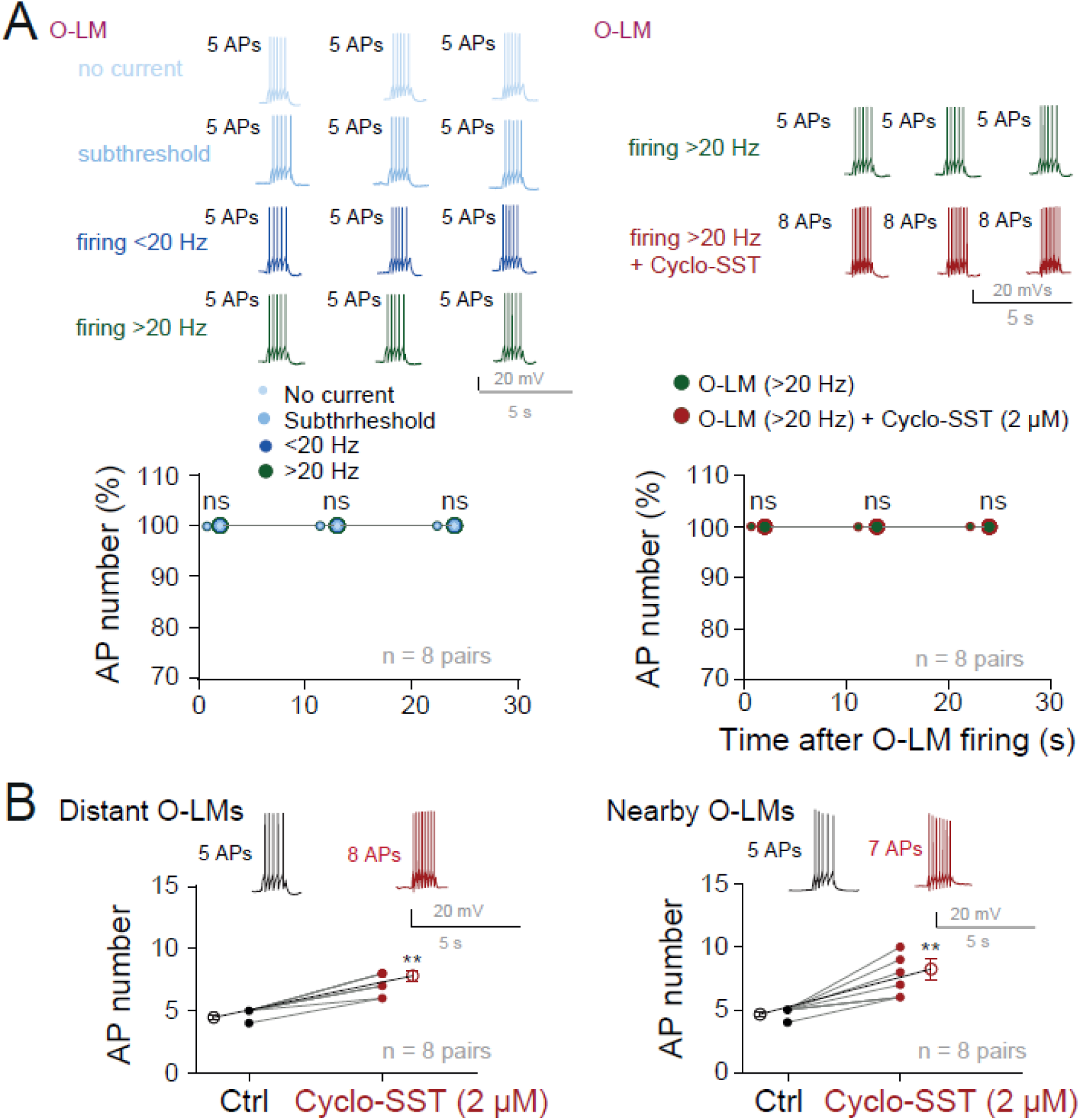
Lack of inhibition at distant O-LM/O-LM cells and enhanced excitability induced by Cyclo-SST in O-LM interneurons. A. Left, firing in the O-LM did not inhibit O-LM cell firing when the two neurons were distant. Right, same experiment in the presence of cyclo-SST (2 µM). ns, Wilcoxon test, not significant. B. Elevated firing in O-LM neurons induced by cyclo-SST when the 2 O-LM cells were nearby (left) or distant (right). **, Wilcoxon test, p<0.01.

**Figure S6.**
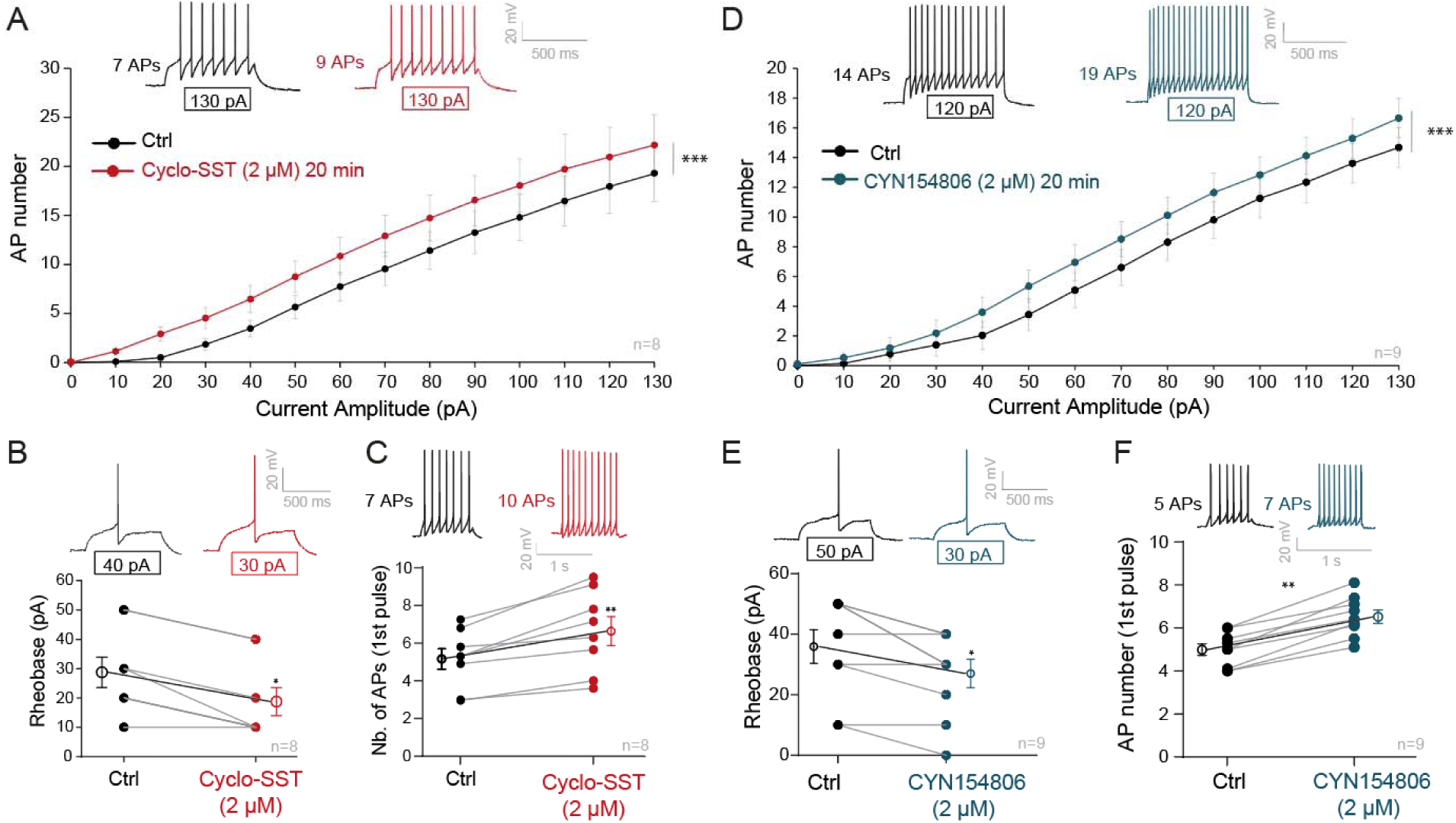
Long-term effect of Cyclo-SST and CYN154806 on intrinsic excitability in O-LM interneurons. A. Input-output curves before and 20 minutes after bath application of the broad SSTR antagonist cyclo-SST. ***, Wilcoxon test p<0.001. B. Reduction of the rheobase induced by prolonged application of cyclo-SST. *, Wilcoxon test p<0.05. C. Enhanced spiking induced by cyclo-SST. **, Wilcoxon test p<0.01. D. Input-output curves before and 20 minutes after CYN154806. ***, Wilcoxon test p<0.001. E. Reduction of the rheobase induced by CYN154806. *, Wilcoxon test p<0.05. F. Enhanced spiking induced by CYN154806. **, Wilcoxon test p<0.01.

**Figure S7.**
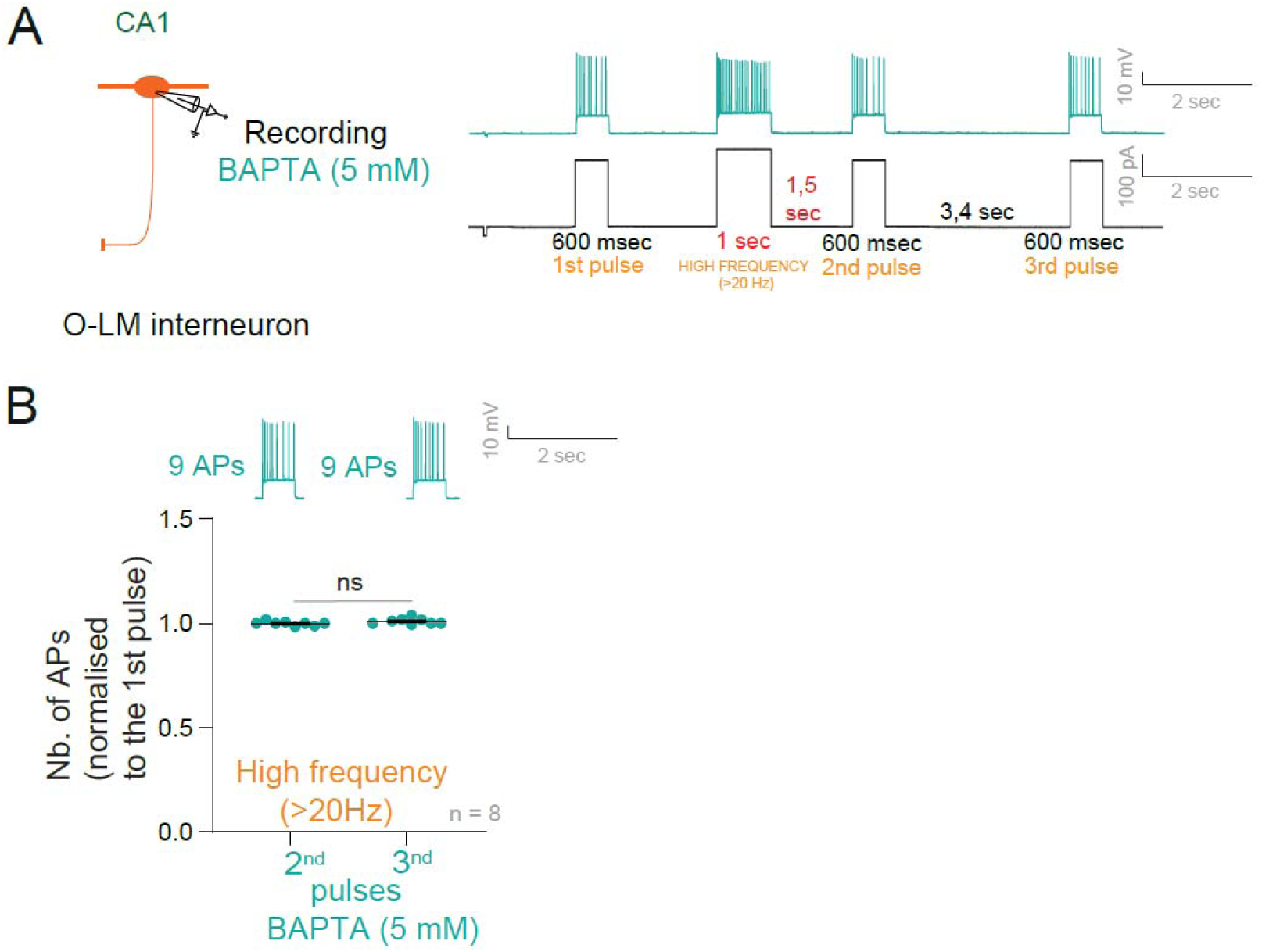
Chelating intracellular calcium in O-LM cells prevents self-inhibition. A. Experimental set-up and protocol. B. Self-inhibition induced by firing at a frequency greater than 20 Hz is blocked when the O-LM is loaded with 5 mM BAPTA. ns, Wilcoxon test, p>0.1.

**Figure S8.**
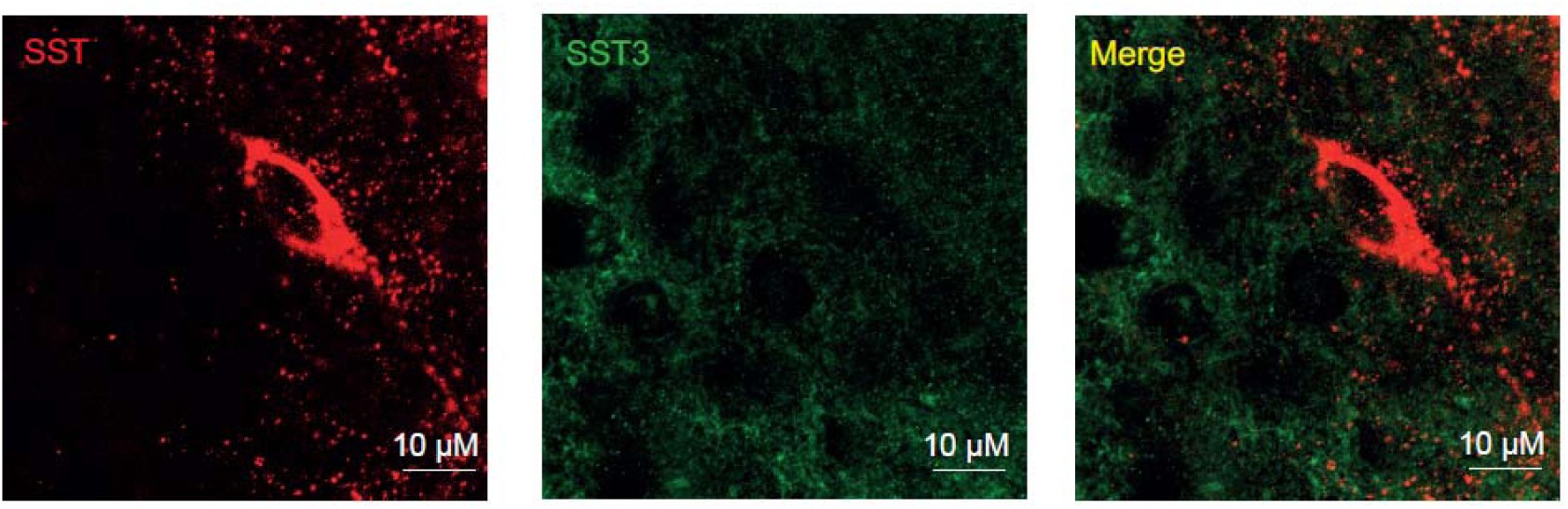
Lack of immunolabeling of SST3 in O-LM interneurons. Left, immunostaining of SST. Middle, SST3 immunostaining. Right, merge image showing the lack of SST3 receptors on O-LM interneuron.

**Figure S9.**
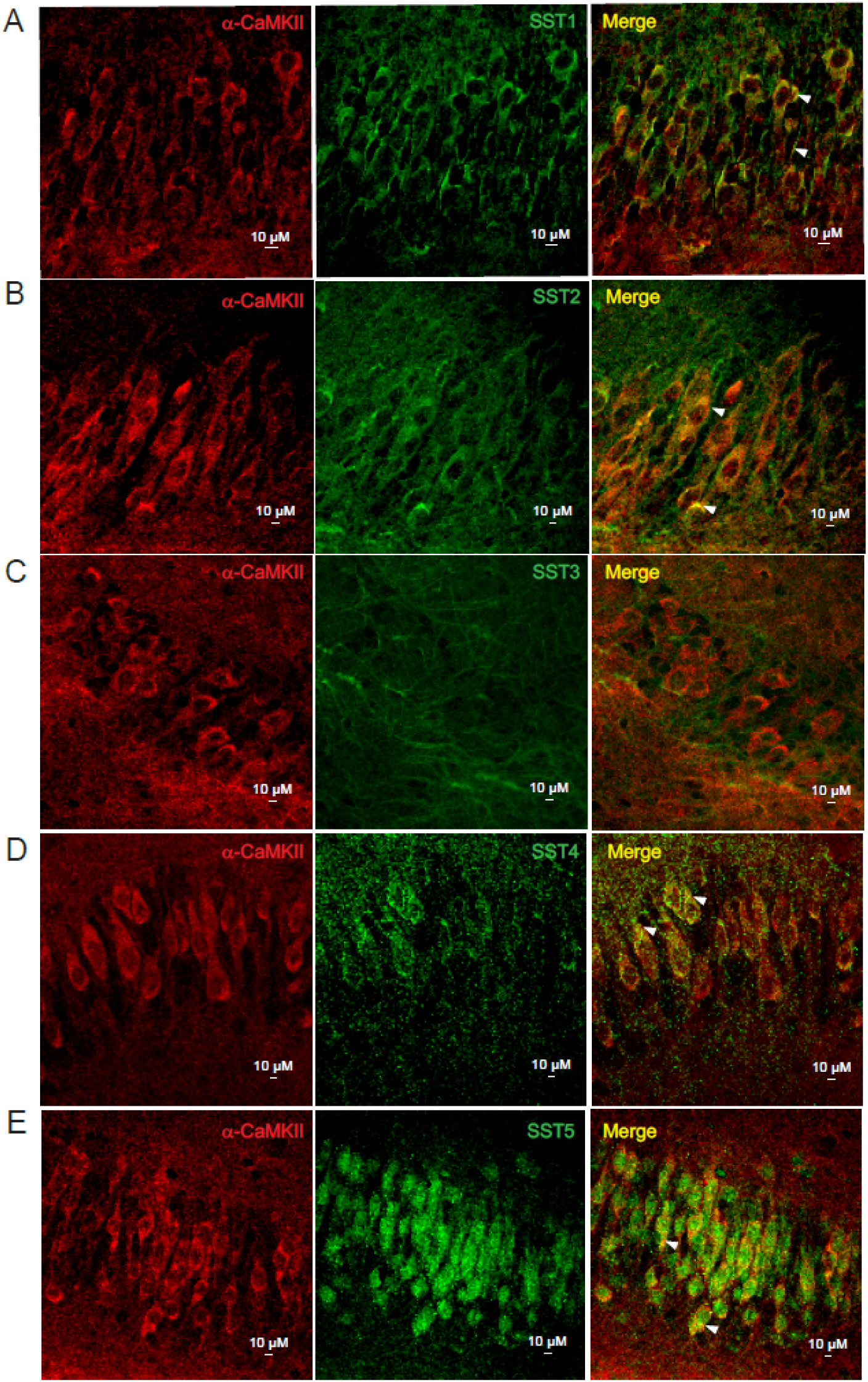
SST receptors in CA1 pyramidal neurons. Immunolabelling of phosphorylated α-CaMKII (left) and SST1 (A, middle), SST2 (B, middle), SST3 (C, middle), SST4 (D, middle) and SST5 (E, middle). Right, merge confocal images showing colocalization in the cell body of phosphorylated α-CaMKII and SST1 (A), SST2 (B), SST4 (D) and SST5 (E) but not with SST3 (C).

**Figure S10.**
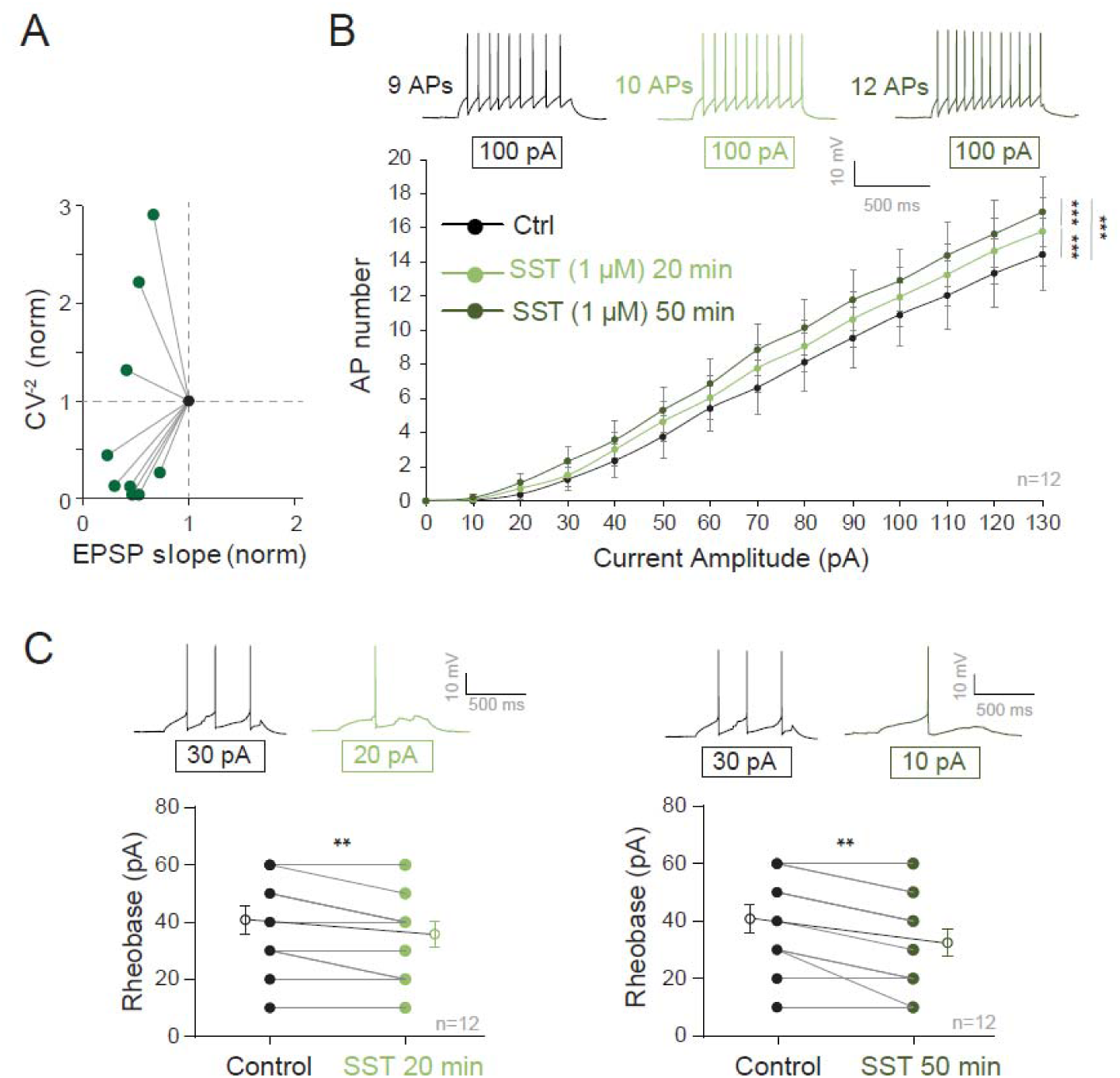
Analysis of CV^-2^ and intrinsic excitability induced by SST. A. Normalized changes in CV^-2^ as a function of EPSP slope changes. B. Intrinsic excitability induced by prolonged application of SST. ***, Wilcoxon test p<0.001. C. Reduction of the rheobase induced by SST for 20 min (left) and 50 min (right)., **, Wilcoxon-test p<0.01.

**Figure S11.**
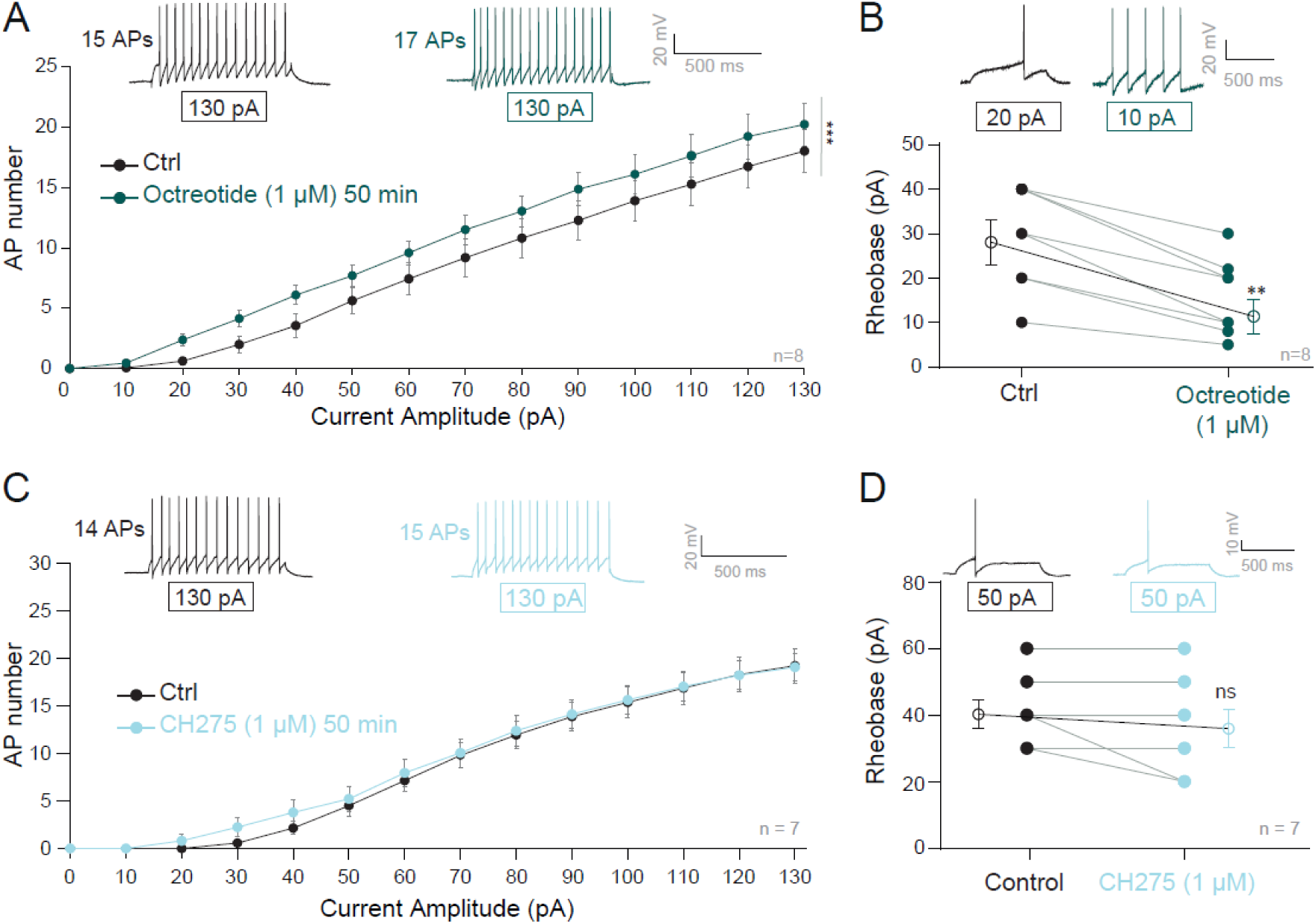
Effects of long-lasting application of the SST2 agonist octreotide and the SST1 agonist CH275 on intrinsic excitability in O-LM interneurons. A. Input-output curves in O-LM interneurons in control and after 50 min in the presence of octreotide (1 µM). ***, Wilcoxon test p<0.001. B. Reduced rheobase in O-LM interneurons. **, Wilcoxon test p<0.01. C. Input-output curves in O-LM interneurons in control and after 50 min in the presence of CH275 (1 µM). D. Rheobase change in O-LM interneuron. ns, Wilcoxon test, not significant.

**Figure S12.**
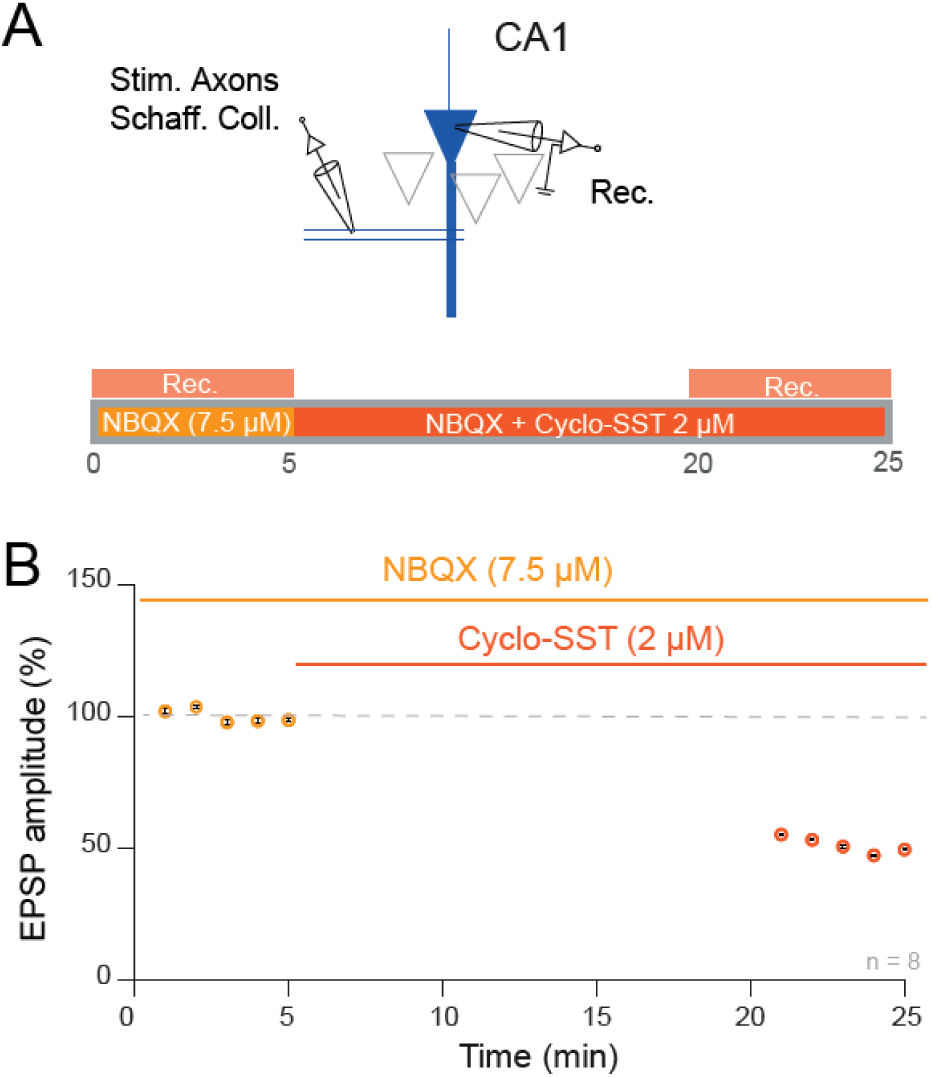
Time-course of the inhibition of NMDA receptor-mediated EPSPs. A. Recording configuration and protocol. B. Time-course of the inhibition of NMDA receptor-mediated EPSP by 2 µM cyclo-SST.

**Figure S13.**
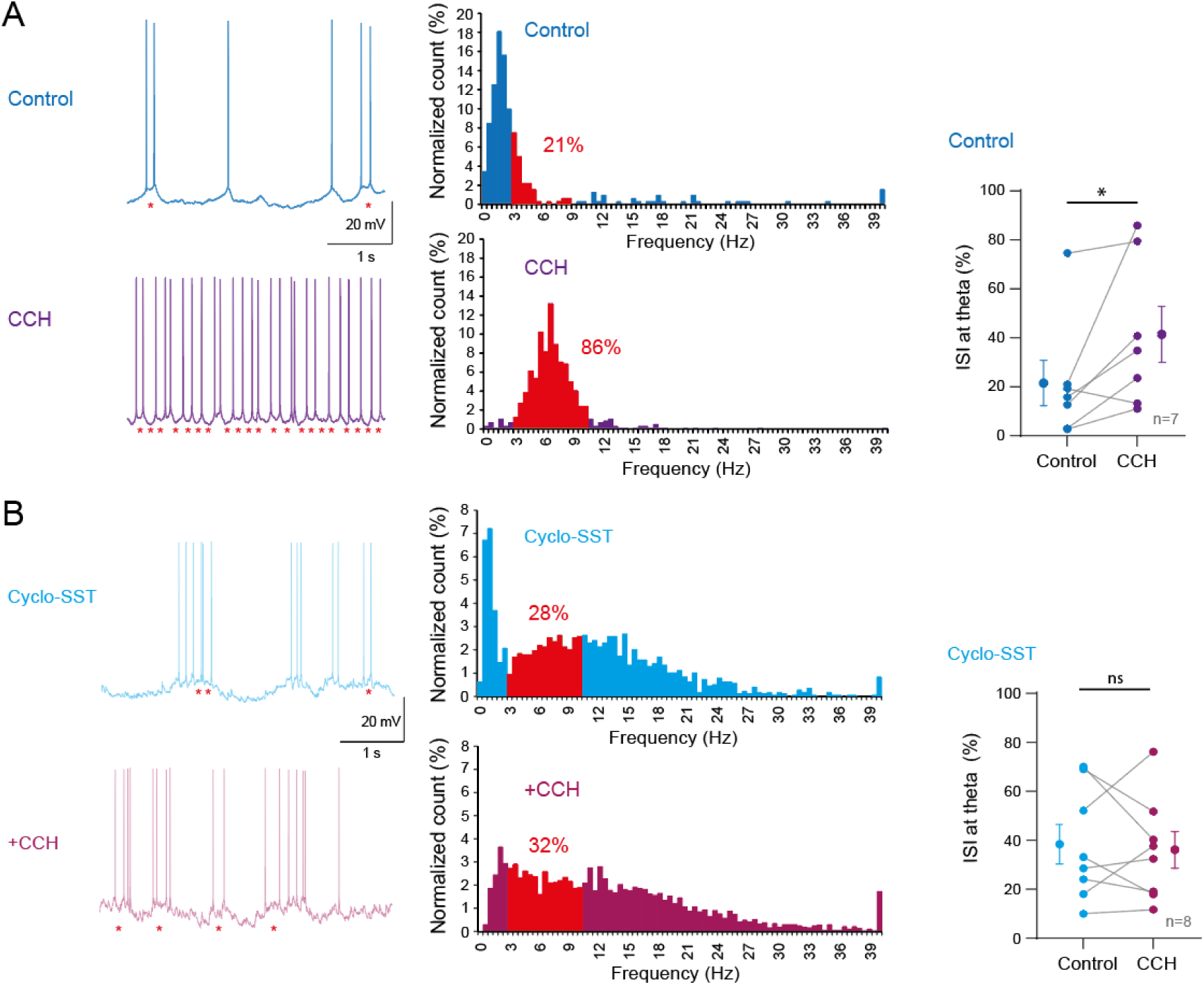
SST receptors blockade prevents the increase of theta firing induced by carbachol (CCH). A. Firing pattern of a CA1 pyramidal cell in control and in the presence of carbachol (CCH). Left, traces; red asterisks indicate inter-spike interval within the theta (θ) frequency range. Note the increase in the number of ISI at θ frequency. Middles, histograms of the instantaneous firing frequency. Note the increase in the proportion of ISI at θ frequency. Right, group data. *, Wilcoxon test p<0.05. B. Firing pattern of a pyramidal cell in the presence of cyclo-SST and in the presence of cyclo-SST + CCH. The number of ISI at θ frequency is about the same in cyclo-SST and in the presence of cyclo-SST and CCH. Middle, histograms. Right, group data. ns, Wilcoxon not significant.

## References

Bezaire MJ, Raikov I, Burk K, Vyas D, Soltesz I (2016) Interneuronal mechanisms of hippocampal theta oscillations in a full-scale model of the rodent CA1 circuit. Elife 5:e18566.

Bi GQ, Poo MM (1998) Synaptic modifications in cultured hippocampal neurons: dependence on spike timing, synaptic strength, and postsynaptic cell type. J Neurosci 18:10464–10472.

Brager DH, Johnston D (2007) Plasticity of intrinsic excitability during long-term depression is mediated through mGluR-dependent changes in I(h) in hippocampal CA1 pyramidal neurons. J Neurosci 27:13926–13937.

Bringuier V, Frégnac Y, Baranyi A, Debanne D, Shulz DE (1997) Synaptic origin and stimulus dependency of neuronal oscillatory activity in the primary visual cortex of the cat. J Physiol 500 ( Pt 3):751–774.

Cammalleri M, Martini D, Timperio AM, Bagnoli P (2009) Functional effects of somatostatin receptor 1 activation on synaptic transmission in the mouse hippocampus. J Neurochem 111:1466– 1477.

Campanac E, Debanne D (2008) Spike timing-dependent plasticity: a learning rule for dendritic integration in rat CA1 pyramidal neurons. J Physiol 586:779–793.

Chamberland S, Grant G, Machold R, Nebet ER, Tian G, Stich J, Hanani M, Kullander K, Tsien RW (2024) Functional specialization of hippocampal somatostatin-expressing interneurons. Proc Natl Acad Sci U S A 121:e2306382121.

Csaba Z, Bernard V, Helboe L, Bluet-Pajot MT, Bloch B, Epelbaum J, Dournaud P (2001) In vivo internalization of the somatostatin sst2A receptor in rat brain: evidence for translocation of cell-surface receptors into the endosomal recycling pathway. Mol Cell Neurosci 17:646–661.

Csaba Z, Dournaud P (2023) Internalization of somatostatin receptors in brain and periphery. Prog Mol Biol Transl Sci 196:43–57.

Debanne D, Boudkkazi S, Campanac E, Cudmore RH, Giraud P, Fronzaroli-Molinieres L, Carlier E, Caillard O (2008) Paired-recordings from synaptically coupled cortical and hippocampal neurons in acute and cultured brain slices. Nat Protoc 3:1559–1568.

Debanne D, Gähwiler BH, Thompson SM (1994) Asynchronous pre- and postsynaptic activity induces associative long-term depression in area CA1 of the rat hippocampus in vitro. Proc Natl Acad Sci U S A 91:1148–1152.

Dournaud P, Gu YZ, Schonbrunn A, Mazella J, Tannenbaum GS, Beaudet A (1996) Localization of the somatostatin receptor SST2A in rat brain using a specific anti-peptide antibody. J Neurosci 16:4468–4478.

Dutton A, Dyball RE (1979) Phasic firing enhances vasopressin release from the rat neurohypophysis. J Physiol 290:433–440.

Extrémet J, Ramirez-Franco J, Fronzaroli-Molinieres L, Boumedine-Guignon N, Ankri N, El Far O, Garrido JJ, Debanne D, Russier M (2023) Rescue of Normal Excitability in LGI1-Deficient Epileptic Neurons. J Neurosci 43:8596–8606.

Fernández-Arroyo B, Jurado S, Lerma J (2025) Understanding OLM interneurons: Characterization, circuitry, and significance in memory and navigation. Neuroscience 578:69–80.

Gasselin C, Inglebert Y, Ankri N, Debanne D (2017) Plasticity of intrinsic excitability during LTD is mediated by bidirectional changes in h-channel activity. Sci Rep 7:14418.

Gastrein P, Campanac E, Gasselin C, Cudmore RH, Bialowas A, Carlier E, Fronzaroli-Molinieres L, Ankri N, Debanne D (2011) The role of hyperpolarization-activated cationic current in spike-time precision and intrinsic resonance in cortical neurons in vitro. J Physiol (Lond) 589:3753–3773.

Gloveli T, Dugladze T, Saha S, Monyer H, Heinemann U, Traub RD, Whittington MA, Buhl EH (2005) Differential involvement of oriens/pyramidale interneurones in hippocampal network oscillations in vitro. J Physiol 562:131–147.

Goldin M, Epsztein J, Jorquera I, Represa A, Ben-Ari Y, Crépel V, Cossart R (2007) Synaptic kainate receptors tune oriens-lacunosum moleculare interneurons to operate at theta frequency. J Neurosci 27:9560–9572.

Gu Z, Alexander GM, Dudek SM, Yakel JL (2017) Hippocampus and Entorhinal Cortex Recruit Cholinergic and NMDA Receptors Separately to Generate Hippocampal Theta Oscillations. Cell Rep 21:3585–3595.

Gustafsson B, Wigström H, Abraham WC, Huang YY (1987) Long-term potentiation in the hippocampus using depolarizing current pulses as the conditioning stimulus to single volley synaptic potentials. J Neurosci 7:774–780.

Hainmueller T, Cazala A, Huang L-W, Bartos M (2024) Subfield-specific interneuron circuits govern the hippocampal response to novelty in male mice. Nat Commun 15:714.

Incontro S, Musella ML, Sammari M, Di Scala C, Fantini J, Debanne D (2025) Lipids shape brain function through ion channel and receptor modulations: physiological mechanisms and clinical perspectives. Physiol Rev 105:137–207.

Incontro S, Sammari M, Azzaz F, Inglebert Y, Ankri N, Russier M, Fantini J, Debanne D (2021) Endocannabinoids Tune Intrinsic Excitability in O-LM Interneurons by Direct Modulation of Postsynaptic Kv7 Channels. J Neurosci 41:9521–9538.

Inglebert Y, Aljadeff J, Brunel N, Debanne D (2020) Synaptic plasticity rules with physiological calcium levels. Proc Natl Acad Sci U S A 117:33639–33648.

Katona L, Lapray D, Viney TJ, Oulhaj A, Borhegyi Z, Micklem BR, Klausberger T, Somogyi P (2014) Sleep and movement differentiates actions of two types of somatostatin-expressing GABAergic interneuron in rat hippocampus. Neuron 82:872–886.

Kazmierska P, Konopacki J (2013) Development of NMDA-induced theta rhythm in hippocampal formation slices. Brain Research Bulletin 98:93–101 Available at: https://www.sciencedirect.com/science/article/pii/S036192301300124X [Accessed June 3, 2025].

Klausberger T, Magill PJ, Márton LF, Roberts JDB, Cobden PM, Buzsáki G, Somogyi P (2003) Brain-state- and cell-type-specific firing of hippocampal interneurons in vivo. Nature 421:844–848.

Klausberger T, Somogyi P (2008) Neuronal diversity and temporal dynamics: the unity of hippocampal circuit operations. Science 321:53–57.

Kreutzberger AJB, Kiessling V, Stroupe C, Liang B, Preobraschenski J, Ganzella M, Kreutzberger MAB, Nakamoto R, Jahn R, Castle JD, Tamm LK (2019) In vitro fusion of single synaptic and dense core vesicles reproduces key physiological properties. Nat Commun 10:3904.

Lawrence JJ, Saraga F, Churchill JF, Statland JM, Travis KE, Skinner FK, McBain CJ (2006) Somatodendritic Kv7/KCNQ/M channels control interspike interval in hippocampal interneurons. J Neurosci 26:12325–12338.

Leão RN, Mikulovic S, Leão KE, Munguba H, Gezelius H, Enjin A, Patra K, Eriksson A, Loew LM, Tort ABL, Kullander K (2012) OLM interneurons differentially modulate CA3 and entorhinal inputs to hippocampal CA1 neurons. Nat Neurosci 15:1524–1530.

Lelouvier B, Tamagno G, Kaindl AM, Roland A, Lelievre V, Le Verche V, Loudes C, Gressens P, Faivre-Baumann A, Lenkei Z, Dournaud P (2008) Dynamics of somatostatin type 2A receptor cargoes in living hippocampal neurons. J Neurosci 28:4336–4349.

Maccaferri G, Roberts JD, Szucs P, Cottingham CA, Somogyi P (2000) Cell surface domain specific postsynaptic currents evoked by identified GABAergic neurones in rat hippocampus in vitro. J Physiol 524 Pt 1:91–116.

Mikulovic S, Restrepo CE, Siwani S, Bauer P, Pupe S, Tort ABL, Kullander K, Leão RN (2018) Ventral hippocampal OLM cells control type 2 theta oscillations and response to predator odor. Nat Commun 9:3638.

Moore SD, Madamba SG, Joëls M, Siggins GR (1988) Somatostatin augments the M-current in hippocampal neurons. Science 239:278–280.

Mulkey RM, Malenka RC (1992) Mechanisms underlying induction of homosynaptic long-term depression in area CA1 of the hippocampus. Neuron 9:967–975.

Müller C, Remy S (2014) Dendritic inhibition mediated by O-LM and bistratified interneurons in the hippocampus. Front Synaptic Neurosci 6:23.

Park E, Mosso MB, Barth AL (2025) Neocortical somatostatin neuron diversity in cognition and learning. Trends Neurosci 48:140–155.

Pittaluga A, Bonfanti A, Raiteri M (2000) Somatostatin potentiates NMDA receptor function via activation of InsP3 receptors and PKC leading to removal of the Mg2+ block without depolarization. Br J Pharmacol 130:557–566.

Pittaluga A, Roggeri A, Vallarino G, Olivero G (2021) Somatostatin, a Presynaptic Modulator of Glutamatergic Signal in the Central Nervous System. Int J Mol Sci 22:5864.

Qiu C, Zeyda T, Johnson B, Hochgeschwender U, de Lecea L, Tallent MK (2008) Somatostatin receptor subtype 4 couples to the M-current to regulate seizures. J Neurosci 28:3567–3576.

Sammari M, Inglebert Y, Ankri N, Russier M, Incontro S, Debanne D (2022) Theta patterns of stimulation induce synaptic and intrinsic potentiation in O-LM interneurons. Proc Natl Acad Sci U S A 119:e2205264119.

Schmid LC, Mittag M, Poll S, Steffen J, Wagner J, Geis H-R, Schwarz I, Schmidt B, Schwarz MK, Remy S, Fuhrmann M (2016) Dysfunction of Somatostatin-Positive Interneurons Associated with Memory Deficits in an Alzheimer’s Disease Model. Neuron 92:114–125.

Schweitzer P, Madamba S, Champagnat J, Siggins GR (1993) Somatostatin inhibition of hippocampal CA1 pyramidal neurons: mediation by arachidonic acid and its metabolites. J Neurosci 13:2033–2049.

Schweitzer P, Madamba S, Siggins GR (1990) Arachidonic acid metabolites as mediators of somatostatin-induced increase of neuronal M-current. Nature 346:464–467.

Schweitzer P, Madamba SG, Siggins GR (1998) Somatostatin increases a voltage-insensitive K+ conductance in rat CA1 hippocampal neurons. J Neurophysiol 79:1230–1238.

Siwani S, França ASC, Mikulovic S, Reis A, Hilscher MM, Edwards SJ, Leão RN, Tort ABL, Kullander K (2018) OLMα2 Cells Bidirectionally Modulate Learning. Neuron 99:404–412.e3.

Song Y-H, Yoon J, Lee S-H (2021) The role of neuropeptide somatostatin in the brain and its application in treating neurological disorders. Exp Mol Med 53:328–338.

Tallent MK, Siggins GR (1997) Somatostatin depresses excitatory but not inhibitory neurotransmission in rat CA1 hippocampus. J Neurophysiol 78:3008–3018.

Tasic B et al. (2018) Shared and distinct transcriptomic cell types across neocortical areas. Nature 563:72–78.

Taxidis J, Madruga B, Safaryan K, Dorian CC, Melin MD, Day Z, Lin MZ, Golshani P (2025) Voltage imaging reveals hippocampal inhibitory dynamics shaping pyramidal memory-encoding sequences. Nat Neurosci.

Tort ABL, Rotstein HG, Dugladze T, Gloveli T, Kopell NJ (2007) On the formation of gamma-coherent cell assemblies by oriens lacunosum-moleculare interneurons in the hippocampus. Proc Natl Acad Sci U S A 104:13490–13495.

Udakis M, Pedrosa V, Chamberlain SEL, Clopath C, Mellor JR (2020) Interneuron-specific plasticity at parvalbumin and somatostatin inhibitory synapses onto CA1 pyramidal neurons shapes hippocampal output. Nat Commun 11:4395.

Varga C, Golshani P, Soltesz I (2012) Frequency-invariant temporal ordering of interneuronal discharges during hippocampal oscillations in awake mice. Proc Natl Acad Sci U S A 109:E2726–2734.

Whim MD, Moss GWJ (2001) A Novel Technique that Measures Peptide Secretion on a Millisecond Timescale Reveals Rapid Changes in Release. Neuron 30:37–50.

